# Origin, specification and differentiation of a rare supporting-like lineage in the developing mouse gonad

**DOI:** 10.1101/2021.09.15.460431

**Authors:** Chloé Mayère, Violaine Regard, Aitana Perea-Gomez, Corey Bunce, Yasmine Neirijnck, Cyril Djari, Pauline Sararols, Richard Reeves, Simon Greenaway, Michelle Simon, Pam Siggers, Diana Condrea, Françoise Kühne, Ivana Gantar, Furong Tang, Isabelle Stévant, Laura Batti, Norbert B. Ghyselinck, Dagmar Wilhelm, Andy Greenfield, Blanche Capel, Marie-Christine Chaboissier, Serge Nef

## Abstract

Gonadal sex determination represents a unique model for studying cell fate decisions. However, a complete understanding of the different cell lineages forming the developing testis and ovary remains elusive. Here, we investigated the origin, specification and subsequent sex-specific differentiation of a previously uncharacterized population of supporting-like cells (SLC) in the developing mouse gonads. The SLC lineage is closely related to the coelomic epithelium and specified as early as E10.5, making it the first somatic lineage to be specified in the bipotential gonad. SLC progenitors are localized within the genital ridge at the interface with the mesonephros and initially co-express *Wnt4* and *Sox9*. SLCs become sexually dimorphic around E12.5, progressively acquire a Sertoli- or granulosa-like identity and contribute to the formation of the rete testis and rete ovarii. Finally, we found that WNT4 is a crucial regulator of the SLC lineage and is required for the formation of the rete testis.

**Teaser:** Description of an uncharacterized multipotent gonadal cell lineage involved in testis and ovary development

## Introduction

Gonadal sex determination is the process by which the bipotential gonad commits to either the testicular or ovarian fate. In the mouse, the gonadal primordium is established at embryonic day 10 (E10), and develops from the thickening of the coelomic epithelium (CE) on the ventromedial surface of the mesonephros (for a review see (*1*)). At this point, CE cells have already acquired a molecular signature unique to the gonads, including the GATA binding protein 4 (*Gata4*), Wilms tumor 1 (*Wt1*), and nuclear receptor steroidogenic factor 1 (*Nr5a1* or *Sf1*) (*2–4*). The genital ridge grows through the proliferation of CE cells that either delaminate and invade the underlying mesenchyme as a result of basement membrane disintegration or as a result of an oriented division pattern, so that daughter cells end up within the developing genital ridge (*2*). Both *in vitro* and *in vivo* lineage tracing experiments and single cell RNA-seq analyses have revealed that the ingressing CE cells constitute the most important source of gonadal somatic cells in both sexes, and contribute to both the supporting and the steroidogenic cell lineages (*5–8*). Initially, these XX and XY CE-derived *Nr5a1*^+^ multipotent progenitor cells do not exhibit obvious sexual dimorphism. By E11.5, a subset of these progenitors adopt a supporting cell fate, before initiating robust sex-dependent genetic programs. This leads to the differentiation of this CE-derived supporting lineage into Sertoli and granulosa cells in the male and female, respectively (*7–9*). In parallel, another subset of these cells maintains its multipotent progenitor state and, starting around E12.5 in XY and E13.5 in XX gonads, acquires a steroidogenic progenitor identity at the origin of Leydig cells and peritubular myoid cells in the developing testis (*10–18*).

A complete characterization of the different cell lineages forming the bipotential gonad is essential for our understanding of the process of sex determination. In particular, it remains unclear whether CE-derived progenitor cells contribute to other uncharacterized populations of the developing testis and ovary. Recent investigations leveraging transcriptomics, immunofluorescence and 3D reconstruction revealed that rete testis precursors possess a dual molecular signature reflecting both a Sertoli (*Sox9*, *Gata4*, *Wt1*, *Nr5a1*) and a rete precursor (*Pax8*, *Krt8* and *Cdh1*) identity, suggesting a common origin with the CE-derived supporting lineage(*19–22*). The rete testis forms an essential bridging system connecting the seminiferous tubules with efferent ducts. In the adult testis, it forms an anastomosing network of epithelium-lined tubules located at testicular mediastinum (*20*). It is connected with, and receives the luminal contents of, the seminiferous tubules. These fluids move from the rete into the efferent ducts and from there into the epididymis and further into the vas deferens. The rete ovarii is the female homolog of the rete testis and is characterized by a set of anastomosed tubules located in the hilus of the ovary that may also extend through the ovarian medulla (*23, 24*). The physiological role of the rete ovarii remains unclear, but it has been proposed to be necessary for the initiation of meiosis and a source of granulosa cells during embryogenesis (*25–27*). Although rete testis and rete ovarii are essential structures connecting the gonads to the reproductive tract, some important gaps remain in the characterization of rete progenitors, their embryonic origin, transcriptomic signature, specification and sex-specific differentiation.

In the present study, we built a transcriptomic atlas of the entire cell population of the developing female and male gonads during the process of mouse sex determination. We identified a previously uncharacterized population of supporting-like cells (SLC) and shed light on the origin and the developmental trajectory of this important cell lineage during the process of testis and ovary development using a combination of single cell RNA sequencing (scRNA-seq) analyses, immunofluorescence, gene expression analysis, *in vivo* cell lineage tracing and loss-of-function experiments.

## Results

### Single-cell transcriptional atlas of gonadal sex determination in mice

We generated a single-cell transcriptomic atlas of mouse XX and XY gonads covering the entire process of sex determination and differentiation, from the emergence of the genital ridges at E10.5, to the late fetal gonads at E16.5, using droplet-based 3’ end scRNA-seq. We collected and sequenced the gonadal cells at five different developmental stages (E10.5, E11.5, E12.5, E13.5, and E16.5) for a total of 94,705 cells after quality controls (**Figure 1A, B** and **Material & Methods**). We found that XX and XY cells tend to overlap at E10.5 and E11.5, as expected for time points prior to gonadal sex determination (**Figure 1C** and **D**). At later stages, cells diverge and display sex-specific gene expression patterns, reflecting the emergence of differentiated cell populations. Gonadal cells were further clustered based on transcriptional similarity using the Leiden algorithm (*28*) (**Figure 1E**). The clustering identified 43 transcriptionally distinct clusters covering XX and XY cells at different stages of gonadal differentiation. Using enriched marker gene expression, we were able to assign a cell type to each of the 43 clusters (**Figure 1E, F** and **Figure S1**). The relative abundance of each cell type was then determined for both the genital ridges and mesonephros at E10.5 and E11.5 and for the testes and ovaries at E12.5, E13.5 and E16.5 (**Figure 1G** and **Table S1**). All major cell populations of the gonad have been identified such as germ cells, Sertoli and granulosa cells, endothelial cells and fetal Leydig cells. It is interesting to note the high proportion of interstitial progenitors in the developing testis and ovary (25% and 58% respectively at E16.5) and the comparatively low abundance of germ cells in the testis (6%) compared to the ovary (34%) at E16.5 as a consequence of mitotic arrest of XY germ cells.

**Figure 1.**
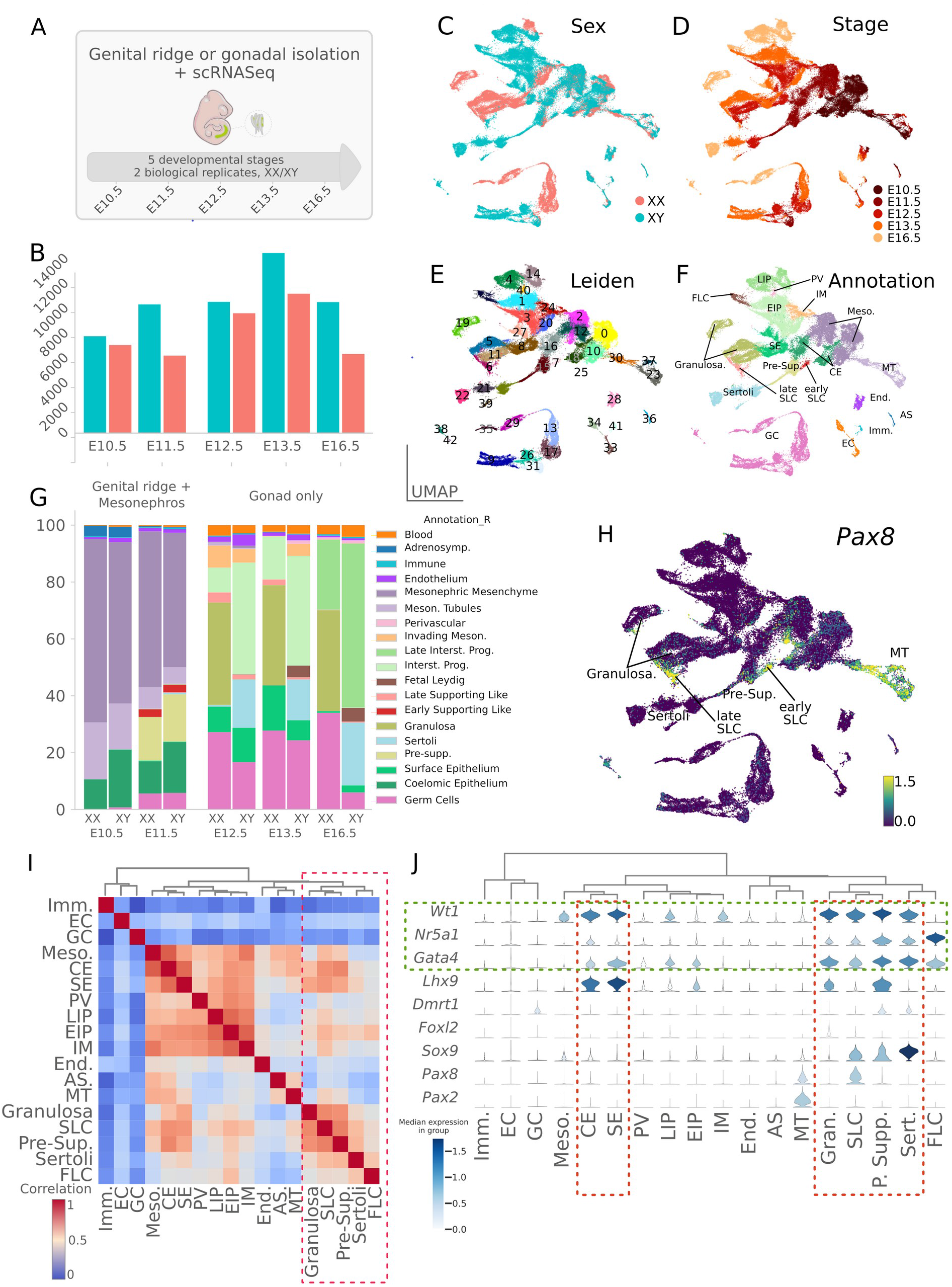
A single-cell atlas of gonadal development and sex differentiation. (**A**) Schematic overview of the experimental procedure and dataset. A representative embryo at E11.5 is shown with (in inset) the region containing the gonads and mesonephros. (**B**) Barplots show the number of single cells RNA sequenced per stage and sex. (**C-F**) UMAP representation of the 94,705 cells colored by sex (**B**), stage (**C**), Leiden clustering (**D**) and annotation of the different cell clusters (**E**). (**G**) Proportion of cell types across sex and developmental stages. (**H**) UMAP representation of the expression level of *Pax8* gene. (**I**) Stacked violin plots showing expression (log-normalized counts) of major somatic cell markers of the gonad in annotated cell populations. Populations were ordered using hierarchical clustering based on correlation (Spearman) distance of expression levels between cell populations. (**J**) Heatmap showing correlation between transcriptomes of the different cell populations. Populations were ordered as in (**I**). Abbreviations: CE, coelomic epithelial cells; SE, surface epithelial cells; Pre-Sup., pre-supporting cells; Sertoli, Sertoli cells; Granulosa, granulosa cells; SLC, supporting-like cells; EIP, early interstitial progenitors; LIP, late interstitial progenitors; FLC, fetal Leydig cells; IM, invading mesonephric cells; PV, perivascular cells, Imm., immune cells; Meso., mesonephric mesenchymal cells; MT, mesonephric tubules; End., endothelial cells; AS, adrenosympathic cells; GC, germ cells; EC, erythrocytes.

### Identification of a previously uncharacterized population of supporting-like cells

Our single-cell transcriptomic analysis also identified all major cell populations derived from the *Nr5a1* lineage, such as supporting progenitors, fetal Leydig, Sertoli, and granulosa cells, but also two clusters containing a previously uncharacterized population of supporting cells, which we refer to as supporting-like cells (SLC). These cells are present in XX and XY gonads from E11.5 to E16.5 (**Figure 1F**). In total, we identified 1,352 unique SLCs forming an early group consisting of 460 cells at E11.5 (hereafter referred to as early SLCs) and 892 cells of later stages (E12.5-E16.5, referred to as late SLCs) both strongly expressing the transcription factor *Pax8* (**Figure 1H**). Although they are part of the same cell population (see below), SLCs group into two separate clusters that reflect two distinct differentiation stages. We identified SLCs in both developing ovaries and testes, with 289 XY and 171 XX early SLCs and 312 XY and 580 XX late SLCs. We observed that the relative abundance of early and late SLCs decreases over time compared to other gonadal cell populations, from 2.7% at E11.5 to 0.1-0.3% at E16.5 (**Figure 1G** and **Table S1**). This is probably due to the significant increase in the number of cells from other cell types during testicular and ovarian development, rather than a decrease in the total number of SLCs. Spearman’s correlation analysis and hierarchical clustering revealed that the transcriptome of the early and late SLC populations share similarities with those of other cell types of the supporting lineage, such as granulosa, pre-supporting and Sertoli cells (**Figure 1I**). SLCs express CE-derived somatic cell markers such as *Nr5a1*, *Wt1*, and *Gata4*, but also *Pax8*, a marker of cells of the mesonephric tubules, the rete testis and rete ovarii (*21, 29*). In contrast, SLCs do not express *Pax2,* a well-known marker of mesonephric tubules (*30*) (**Figure 1J**). Overall, the expression profile of SLCs suggests that they are closely related to the CE-derived supporting cells and potentially involved in the formation of the rete ovarii in females, and the rete testis and its connection to efferent ducts in males.

### XY SLCs share a common CE-derived origin but diverge from the Sertoli lineage before the time of sex determination

To characterize the specification and fate of SLCs in the developing testis, we selected all of the 11 XY cell clusters expressing the three marker genes present in CE-derived gonadal cells (*Nr5a1*, *Gata4* and *Wt1*) between E10.5 and E16.5 (clusters #6, 7, 8, 15, 16, 18, 21, 22, 25, 27 and 39, see **Figure 1E**). We then used the PAGA algorithm (Plass et al., 2018; Wolf et al., 2019) to generate a consolidated lineage graph that included all selected cell types rooted to the E10.5 CE cluster (**Figure 2A**). The reconstruction of the CE-derived cell lineages in the developing XY gonad revealed three distinct developmental trajectories, with all cells emerging initially from the CE-derived progenitor cells at E10.5 and E11.5 (clusters #16 and #18, **Figure 2A-C**). Three different cell lineages emerge from these CE-derived progenitors: the gonadal surface epithelium (SE) that expresses the *Upk3b* marker gene (clusters #8 and #27, **Figure 2A-E**); the supporting cell lineage expressing sequentially *Sry*, *Sox9* and *Amh* during the process of Sertoli cell differentiation (clusters #15 then #25, #21, #22 and #39, **Figure 2A-D** and **F-H**); and the SLC lineage, expressing *Pax8* as well as *Sox9*, but not *Sry* (see clusters #7 and #6, **Figure 2A-D, G** and **I**). The *Pax8* gene appears also transiently expressed in 5% of XY pre-supporting cells at E11.5 (threshold defined as 0.5 log normalized counts see also **Figure 2I**) suggesting that its expression is not completely SLC-specific. Cell cycle analysis using scRNA-seq data (*31*) revealed that cells in the coelomic epithelium, surface epithelium, and Sertoli cells actively proliferate, whereas pre-supporting cells expressing *Sry* and SLCs appear to remain in a quiescent state (**Figure 2J).** The absence of proliferation in the SLC lineage was confirmed by whole-mount immunofluorescence of gonads pulsed with 5-Ethynyl-2’-deoxyuridine (EdU) incorporation. Absence of EdU labelling of PAX8^+^/GATA4^+^ cells confirms that SLCs arrest cycling during the process of sex determination at E11.5 and early testis development at E13.5 (**Figure S2**).

**Figure 2.**
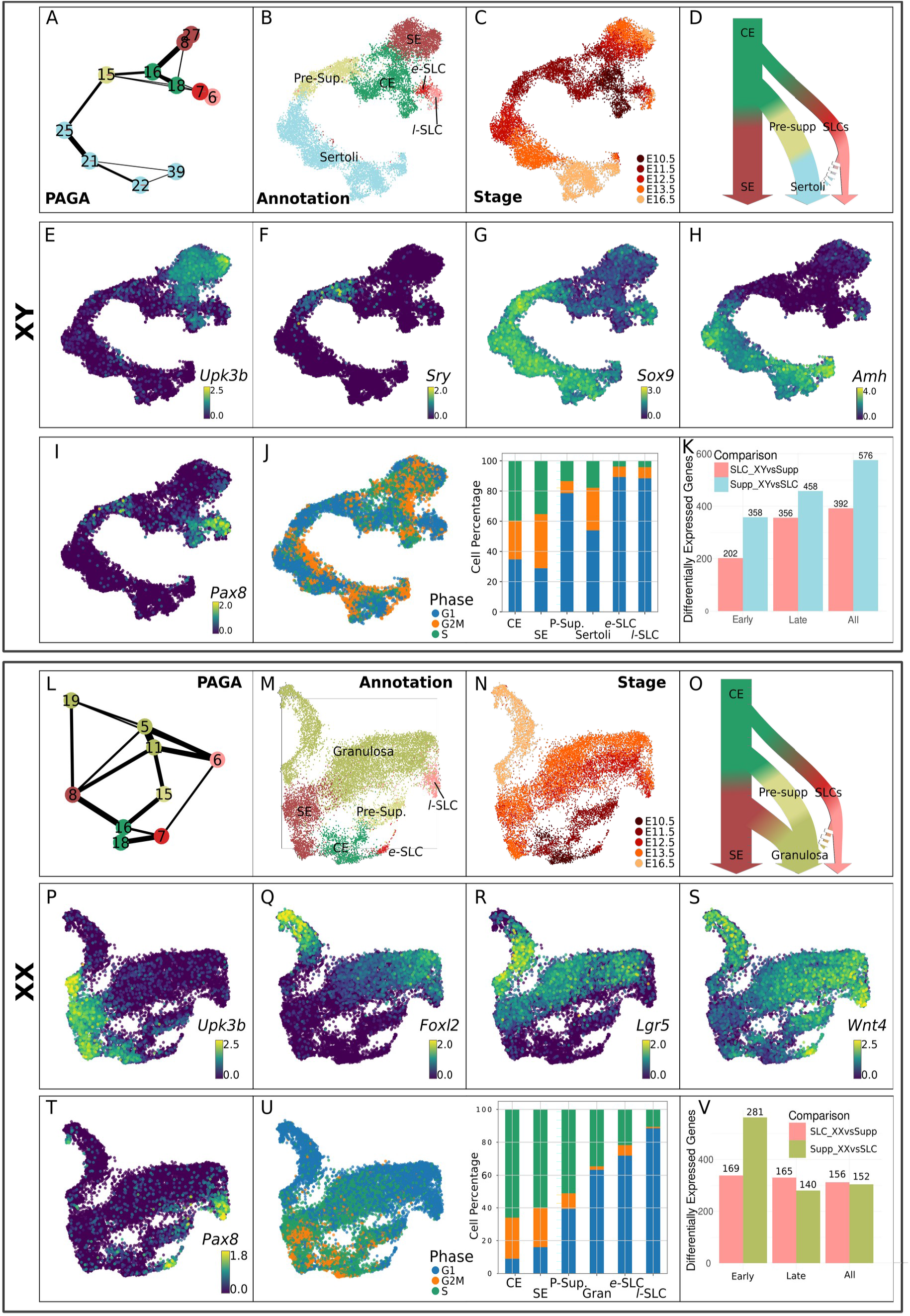
Characterization of XY and XX supporting-like cell (SLC) populations. The analysis was performed with both XY (**A-K**) and XX (**L-V**) gonadal cells. PAGA representation of the clusters of XY (**A**) and XX (**L**) cells expressing *Nr5a1*, *Gata4* and *Wt1*. Each node is a Leiden cluster and the links between clusters represent confidence of relation between two clusters. (**B, C, M, N**) UMAP projection of gonadal cell lineage at the origin of the supporting cells and SLCs, colored by cell annotation (**B, M**) and by developmental stage (**C, M**). (**D**) Simplified model of the cell lineage reconstruction in the developing testis (**D**) and ovary (**O**). UMAP representations of XY (**E-I**) and XX (**P-T**) cells colored by expression levels (log-normalized counts) of selected genes. UMAP and barplot representation of the cell cycle phase of XY (**J**) and XX (**U**) cells. Barplot illustrates the proportion of cells in the different cell cycle phase for each population. (**K, V**) Barplots representing the number of differentially-expressed genes between SLCs and the supporting lineage either in XY (**K**) or XX (**V**) developing gonads at early (E11.5) and late (E12.5, E13.5 and E16.5) stages. Abbreviations: CE, coelomic epithelial cells; SE, surface epithelial cells; P-Sup., pre-supporting cells; Sertoli, Sertoli cells; Granulosa; *e*-SLC, early supporting-like cells; *l*-SLC, late supporting-like cells.

To investigate the differences between SLCs and the supporting lineage in the developing testis, we performed differential expression analysis (*32*). As the number of SLCs decreases with time, we grouped these cells into two groups to maintain sufficient statistical power for the analysis: early SLCs (cells at E11.5) and late SLCs (cells between E12.5 and E16.5). We observed a large number of differentially expressed genes (DEGs) both at E11.5 (560 genes) and at later stages (814 genes) (**Figure 2K**). Genes overexpressed in XY SLCs were linked with gene ontology terms such as the “BMP pathway”, “mesonephric tubule development”, and “regulation of TGFβ stimulus”, suggesting that although these cells do not originate from mesonephric tubules, they share some common transcriptomic features. Among the DEGs overexpressed in XY SLCs, we find SLC lineage markers such as *Pax8*, *Ncam1*, *Tbx2* and *Ennp2* but also genes of the WNT/β-catenin signaling pathway such as *Wnt4*, *Lef1*, *Nkd1*, and the transcription factor *Nr0b1* (*Dax1*). Similarly, *Bmp4* and the *Id1, Id2 and Id3* genes, targets of the BMP pathway, are also overexpressed in XY SLCs consistent with the GO term “BMP pathway”. As expected, genes overexpressed in the supporting cell lineage were associated with terms associated with “male gonad development” (see **Data S1** for more details on DEGs and GO biological processes enrichment analysis) and include classical Sertoli cell markers such as *Amh, Ptgds, Tesc, md2, Nedd9, Mro, Dhh, Cst8*.

Overall, our lineage tracing reconstruction is consistent with a coelomic origin of the SLCs that subsequently adopt a developmental trajectory independent of the supporting cell lineage before or around the time of sex determination (**Figure 2D**).

### XX SLCs are already specified by E11.5 and contribute to the pool of granulosa cells

To characterize the specification and fate of SLCs in the developing ovaries, we used the PAGA algorithm to generate a consolidated lineage graph that includes all the nine XX cell clusters between E10.5 and E16.5 expressing *Nr5a1*, *Gata4* and *Wt1* (*i.e.* clusters #5, 6, 7, 8, 11, 15, 16, 18 and 19, see **Figure 1D, F-H** and **Figure 2L**). Due to the existence of distinct populations of granulosa cells, as well as cellular heterogeneity in the precursors of the granulosa cell lineage (*33–35*), the resulting PAGA graph is more complex than its testicular counterpart. Nevertheless, we observed three different cell lineages originating from CE-derived progenitors (clusters #18 and 16, **Figure 2L-O**): the gonadal surface epithelium (SE) cells that expresses the *Upk3b* marker gene (cluster #8, **Figure 2L-O** and **P**); the supporting cell lineage sequentially expressing *Foxl2* and *Lgr5* during the process of granulosa cell differentiation (clusters #15, #11, #5 and #19, **Figure 2L-O, Q** and **R**); and the SLC lineage, expressing *Pax8* and *Enpp2* (clusters #7 and #6, **Figure 2L-O, S** and **T**). As for their male counterpart, the XX SLCs are already specified by E11.5 (cluster #7). In respect of *Pax8* gene, it also appears transiently expressed in 5% of XX pre-supporting cells at E11.5 (threshold defined as 0.5 log normalized counts; see **Table S1** and **Figure 2P**) suggesting that it is not specific to SLCs in the developing gonad. Cell cycle analysis revealed that CE and SE cells are highly proliferative, whereas late SLCs are in a quiescent phase (**Figure 2U**). Interestingly, PAGA analysis showed strong connections between granulosa cells (clusters #11, #5 and #19) and cells from the SE (cluster #8) as well as late SLCs (cluster #6, **Figure 2L**). Similarly, in the UMAP representations, the SE cluster (#8) showed a continuous connection with *Lgr5*-expressing granulosa cells, while the late SLCs were connected to *Foxl2*-expressing granulosa cells (**Figure 2L-N**). It is therefore likely that granulosa cells are derived from multiple sources of progenitors, as recently described (*36*). Our transcriptomic data suggest that granulosa cells initially derived from pre-supporting cells around E12.5 (cluster #15), while at later stages, both SE cells (cluster #8) and late SLCs (cluster #6) may contribute to the granulosa cell pool.

Differential expression analysis between XX SLCs and the supporting lineage revealed that the set of DEGs is significantly smaller compared to XY SLCs and Sertoli cells, both at E11.5 (450 genes) and at later stages (305 genes) (**Figure 2V**). This confirmed that XX and XY SLCs are transcriptionally closer to granulosa cells than to Sertoli cells (compare **Figure 2K** and **V**). Genes overexpressed in XX SLCs were linked with terms such as “response to BMP”, “negative regulation of growth factor stimulus”, “mesonephric tubule development”. It includes SLC marker genes such as *Pax8*, *Ncam1*, *Tbx2* and *Ennp2* but also the pro-testis factor *Sox9* as well as genes of the WNT/β-catenin signaling pathway such as *Wnt4*, *Lef1*, *Nkd1* and the BMP responding genes *Id1, Id2* and *Id3*. As expected genes overexpressed in the granulosa cell lineage were associated with terms such as “female gonad development” and include genes such as *Lhx9*, *Fst*, *Irx3*, *Bmp2*, *Lgr5* (see **Data S2** for more details on DEGs and GO biological processes enrichment analysis).

Overall, our lineage tracing reconstruction suggests that XX SLCs adopt a developmental trajectory independent of the supporting lineage before or around the time of sex determination and may also contribute to the pool of granulosa cells (**Figure 2O**).

### SLCs are specified at E10.5 and become gradually sexually dimorphic from E12.5

Having shown that both XY and XX SLCs adopt a developmental trajectory that is independent from the supporting cell lineage, we next aimed to identify their specification at E10.5. We next examined the emergence of sexual dimorphism in the SLC lineage as the XY and XX SLCs progress through gonadal development. To this end, we used the PAGA algorithm to generate a consolidated lineage graph that includes the CE clusters at E10.5 (cluster #18) and E11.5 (cluster #16), pre-supporting cell cluster at E11.5 (cluster #15), as well as early and late SLC clusters between E11.5 and E16.5 (i.e. clusters #6 and #7, see **Figure 1D, F-H**). Applying a Leiden clustering restricted to CE clusters, we found that the CE-derived progenitors represent a heterogeneous population of cells that can be classified into seven sub-clusters (thereafter named CE-0 to CE-6, **Figure 3A**). At E10.5, only three sub-clusters express the classical CE-derived markers *Gata4* and *Nr5a1*, namely CE-1, CE-5 and CE-6. Interestingly, our PAGA graph indicated that sub-clusters CE-1 and CE-5 are connected to sub-cluster CE-6, which is connected to the early and late SLCs clusters (**Figure 3A**). While cells from sub-clusters CE-1 and CE-5 expressed classical CE-derived markers such as *Gata4*, *Nr5a1*, they were, however, *Pax8* negative, therefore representing the canonical CE-derived progenitors. In contrast, cells from sub-cluster CE-6 expressed not only *Gata4* and *Nr5a1*, but also *Pax8,* suggesting that they are in the process of transitioning toward the SLC lineage (see **Figure 3B, C** and **D**). To better characterize the transcriptomic changes related to the specification of the SLC lineage at E10.5, we then investigated which genes were differentially expressed between the *Pax8*-expressing cells of the CE-6 sub-cluster and the other two sub-clusters which represent the E10.5 CE population (CE-1 and CE-5). For this purpose, we retained only genes with a mean log fold change (LogFC) > 0.25 and an FDR-adjusted P-value <0.05 (**Data S3**). Among the 112 genes differentially expressed, we found 59 genes downregulated in the CE-6 sub-cluster including the CE markers *Lhx9*, *Upk3b* but also genes such as *Gata4, Tcf21* and *Wt1*. In addition, we observed 53 upregulated genes including marker genes of the SLC lineage such as *Pax8*, *Ncam1* and *Enpp2* as well as genes known to be involved in the process of gonadal determination such as *Nr0b1*, *Sox9*, *Wnt4* as well as *Igf1* and *Igf2*. These results indicate that the SLC lineage specification is initiated as early as E10.5 in a subset of CE-derived progenitors with the latter rapidly acquiring SLC transcriptomic markers such as *Pax8, Ncam1* and *Enpp2*.

**Figure 3.**
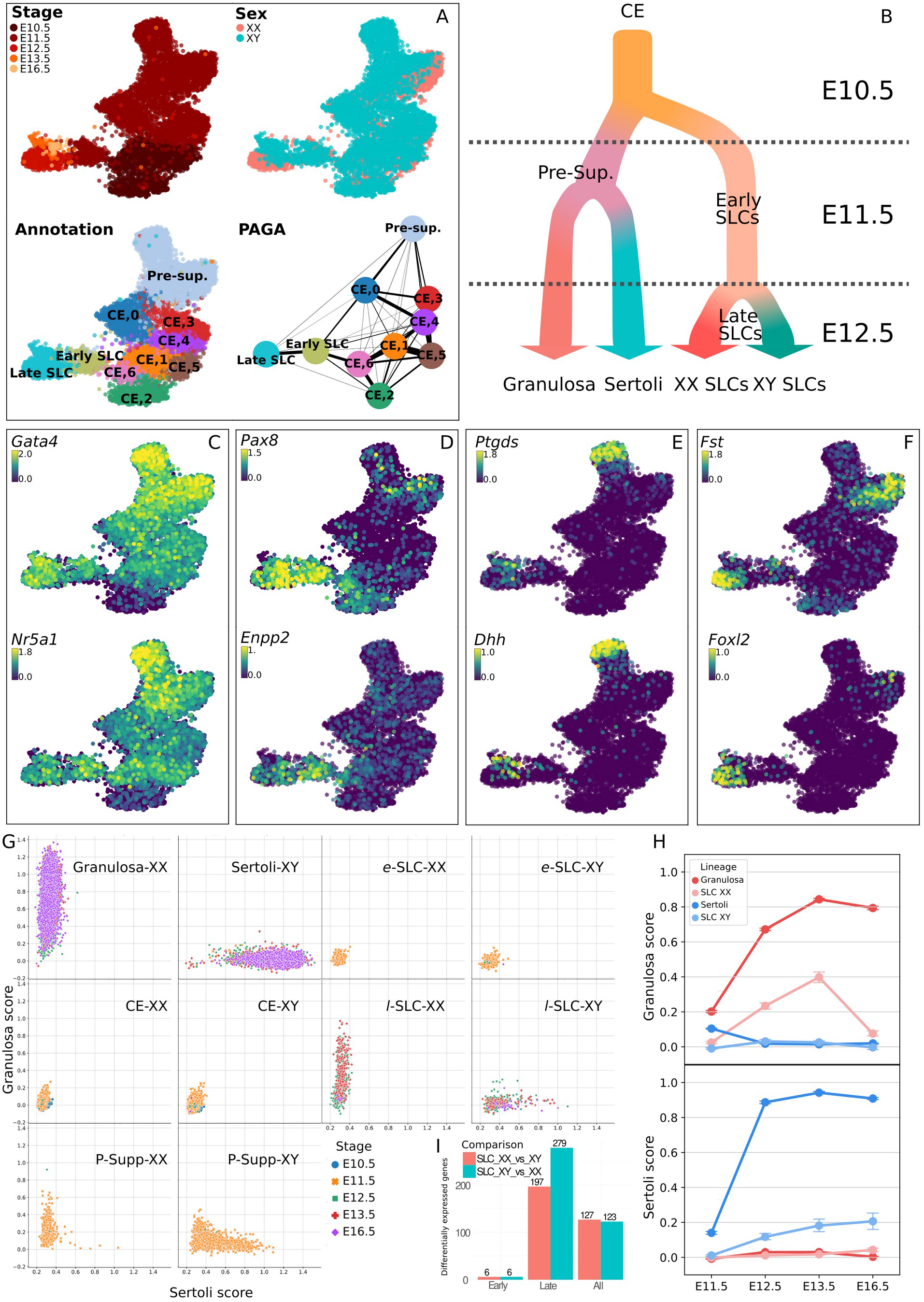
Sex-specific differentiation of XX and XY SLCs. (**A**) UMAP projection of SLC cell lineage colored by developmental stage, genetic sex and annotation with subclustering applied to the CE. The fourth panel shows the PAGA graph representation where each node is a Leiden cluster and the links between clusters represent confidence of relation between two clusters. (**B**) Schematic model of SLC lineage specification. (**C-F**) UMAP representation colored by expression levels (log-normalized counts) of CE-derived markers *Gata4* and *Nr5a1* (**C**), SLC markers *Pax8* and *Enpp2* (**D**), Sertoli cell markers *Ptgds* and *Dhh* (**E**), and granulosa cell markers *Fst* and *Foxl2* (**F**). (**G**) Scatterplots representing granulosa and Sertoli scores of individual cells using a LASSO model. A plot was generated for each cell type and individual cells were colored by developmental stage. (**H**) Pointplots showing granulosa and Sertoli cell scores in XX and XY SLC and supporting cells across time. (**I**) Barplots representing the number of significantly over-expressed genes found when comparing XX to XY SLCs at early (E11.5), late (E12.5, E13.5 and E16.5) or all stages.

Initially, early XX and XY SLCs at E10.5 and E11.5 remain sexually undifferentiated and do not exhibit sexual dimorphism. Sexual dimorphism in SLCs develops gradually from E12.5 onward with the upregulation of Sertoli (e.g. *Ptgds*, *Dhh*) and granulosa (e.g. *Foxl2*, *Fst*) cell-specific markers in late SLCs (**Figure 3E** and **F**). Moreover, XX and XY SLCs at E11.5 co-express the pro-ovarian gene *Wnt4* and the pro-testicular gene *Sox9* at similar levels (**Figure S3A**), suggesting that both pathways may be active in early SLCs. This observation is consistent with whole-mount in situ hybridization (WISH) revealing the presence of *Sox9* and *Wnt4* transcripts in the antero-dorsal part of both E13.5 testis and ovary where the rete testis and rete ovarii are forming (see arrows in **Figure S3B**).

To better characterize how differentiating SLCs acquire their Sertoli or granulosa characteristics, we trained a LASSO model to determine a score based on the molecular signature of both cell types. We have identified a set of 167 and 74 genes that define the molecular signature of Sertoli and granulosa cells, respectively (see **Data S4**). According to this model, XY and XX progenitor cells of the CE exhibit low Sertoli and granulosa scores, consistent with their undifferentiated progenitor status (**Figure 3G-I**). In contrast, and as expected, pre-supporting cells, Sertoli cells and granulosa cells are characterized by increased Sertoli and granulosa scores. Similarly, from E12.5 onward we observed a progressive increase of the Sertoli score for XY SLCs, and a gradual increase of the granulosa score in XX SLC cells. In-depth analysis indicated that late XY SLCs start to gradually express Sertoli-specific genes such as *Dhh*, *Amh*, *Ptgds*, *Aard*, *Tesc*, *Vnn1*, *Nedd9*, *Cbln4*, while maintaining the expression of genes enriched in SLCs such as *Pax8* and *Ncam1* (**Figure S4**). Similarly, late XX SLCs gradually upregulated classical granulosa markers, such as *Wnt4*, *Foxl2*, *Fst*, *Irx3*, and downregulated *Sox9* while maintaining the expression of *Pax8* and *Ncam1*. Although the increase in expression of Sertoli or granulosa markers in late XY and XX LSCs is significant, it does not reach the expression levels present in the supporting cell line (**Figure S4B** and **C**). Overall, these results demonstrate that SLC lineage specification occurs around E10.5 and that the establishment of sexual dimorphism in the SLC lineage starts at E12.5 (**Figure 3B**). Then, differentiating XY and XX SLCs gradually acquire a molecular signature similar to Sertoli and granulosa cells respectively, reinforcing the possibility that SLCs have the capacity to differentiate into either of these cell types.

### SLCs arise in both XY and XX genital ridges at E11.0 and become restricted to either the rete testis or rete ovarii

To confirm the results obtained using scRNA-seq and to characterize the spatiotemporal localization of the SLC population during the process of gonadal development, we analyzed the expression of the SLC marker PAX8 by whole-mount immunofluorescence (IF) of wildtype XX and XY genital ridges from E10.5 to E12.5. Our analysis revealed that a population of cells faintly expressing PAX8 and the gonadal somatic cell marker GATA4 at around E11.0 are present throughout the XX and XY genital ridge at the border between the gonad and the mesonephros (**Figure 4**, XY E11.0). These IF data were consistent with the identification of a small cluster of CE-derived gonadal cells expressing *Pax8* at E10.5 by scRNA-seq (**Figure 3A** and **D**). At early stages, the PAX8^+^ gonad/mesonephros border domain is adjacent to the mesonephric tubules, making it difficult to distinguish SLC PAX8 from mesonephric PAX8. However, PAX8 is generally expressed at higher levels in the mesonephric tubules than in the PAX8^+^/GATA4^+^ SLC presumptive rete progenitors. At E11.0, we also observed rare PAX8^+^/GATA4^+^/SOX9^+^ cells in the anterior part of the XY gonads (arrowhead in **Figure 4**). At E11.5, a portion of the PAX8^+^/GATA4^+^ population also expressed SOX9 in both XY and XX gonads. Similar to Sertoli and granulosa cells, PAX8^+^/GATA4^+^ cells along the border of the gonad were NR2F2^−^. IF analysis of laminin subunit Beta-1 (LAMB1) reveals that PAX8^+^/GATA4^+^ cells along the gonad/mesonephros border are associated with a basement membrane (**Figure 4**, XX E10.5 and XX E12.5). LAMB1 expression is continuous within the PAX8^+^/GATA4^−^ mesonephric tubules and the PAX8^−^/GATA4^+^ coelomic epithelium from E10.5 to E12.5.

**Figure 4.**
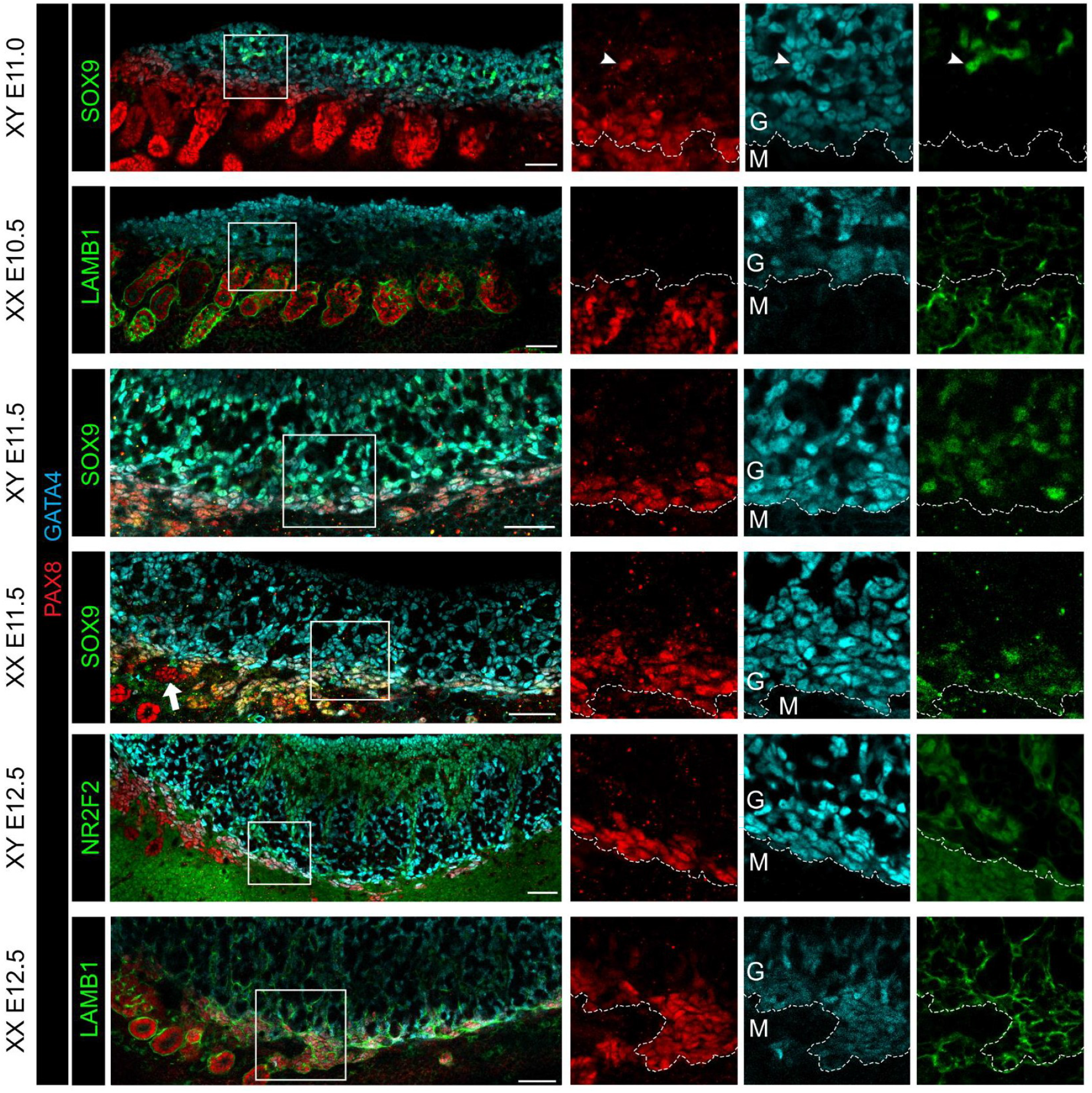
A population of gonadal cells expressing both GATA4 and PAX8 is present from E10.5-E11.0 along the genital ridge next to the mesonephros. Whole-mount immunofluorescence of XY and XX genital ridges from E10.5 to E12.5. Boxes in merged images indicate regions shown as isolated channels on right. The dotted line correspond to the border that separates the GATA4^+^ cells of the genital ridge (G) from the mesonephric cells (M). LAMB1 is expressed in the basement membrane throughout the CE and mesonephric tubules. SOX9 overlaps with a subset of PAX8+/GATA4+ cells in XY and XX tissues at E11.5. NR2F2 is a marker of gonad interstitial cells and mesonephric mesenchyme. Arrowhead indicates SOX9^+^/PAX8^+^ cell within the XY gonad at E11.0. Arrow indicates PAX8^+^/GATA4^−^ cells adjacent in the region of the presumptive rete that are not within mesonephric tubules. Scale bar = 50um.

Overall, the PAX8 expression analysis was consistent with our scRNA-seq analysis. IF further revealed that SLC progenitor cells can be found at the border to the mesonephros in XY and XX genital ridges as early as E11.0 and that they contribute to the rete testis and rete ovarii, respectively.

### *Pax8*^+^ progenitors contribute to both rete testis/ovarii and to the pool of Sertoli and granulosa cells

To determine the fate of SLC progenitors and their spatiotemporal localization in the developing testis and ovary, we first used light sheet fluorescence microscopy on cleared whole gonads to obtain a high-resolution tridimensional visualization of the urogenital system. The technique allowed us to examine the anatomical distribution of the targeted cells in the intact gonad, by preserving the three-dimensional information. The evaluation was carried out at E13.5 when the rete structure develops, at E16.5, and at birth (P0) when testis and ovary development is well underway and rete formation is completed (*19, 37*). To this end, we labelled *Pax8*-expressing cells and all their derivatives as well as *Nr5a1*-expressing cells using a transgenic line *Pax8:Cre;Rosa26:tdTomato;Nr5a1:GFP* (*7, 38, 39*). In the developing testis (**Figure 5A**), RFP^+^ cells were primarily clustered in the cranio-dorsal region of the gonad, corresponding to the rete testis. In addition, a small set of RFP^+^ cells were also observed scattered throughout the testis. Similarly, in the developing ovary (**Figure 5A**), RFP^+^ cells were present along the antero-dorsal region of the gonad near the mesonephros. The clustering in the anterior part is not as pronounced as in the testis. We also observed some RFP^+^ cells scattered within the developing ovary, although to a lesser extent than the male counterpart.

**Figure 5.**
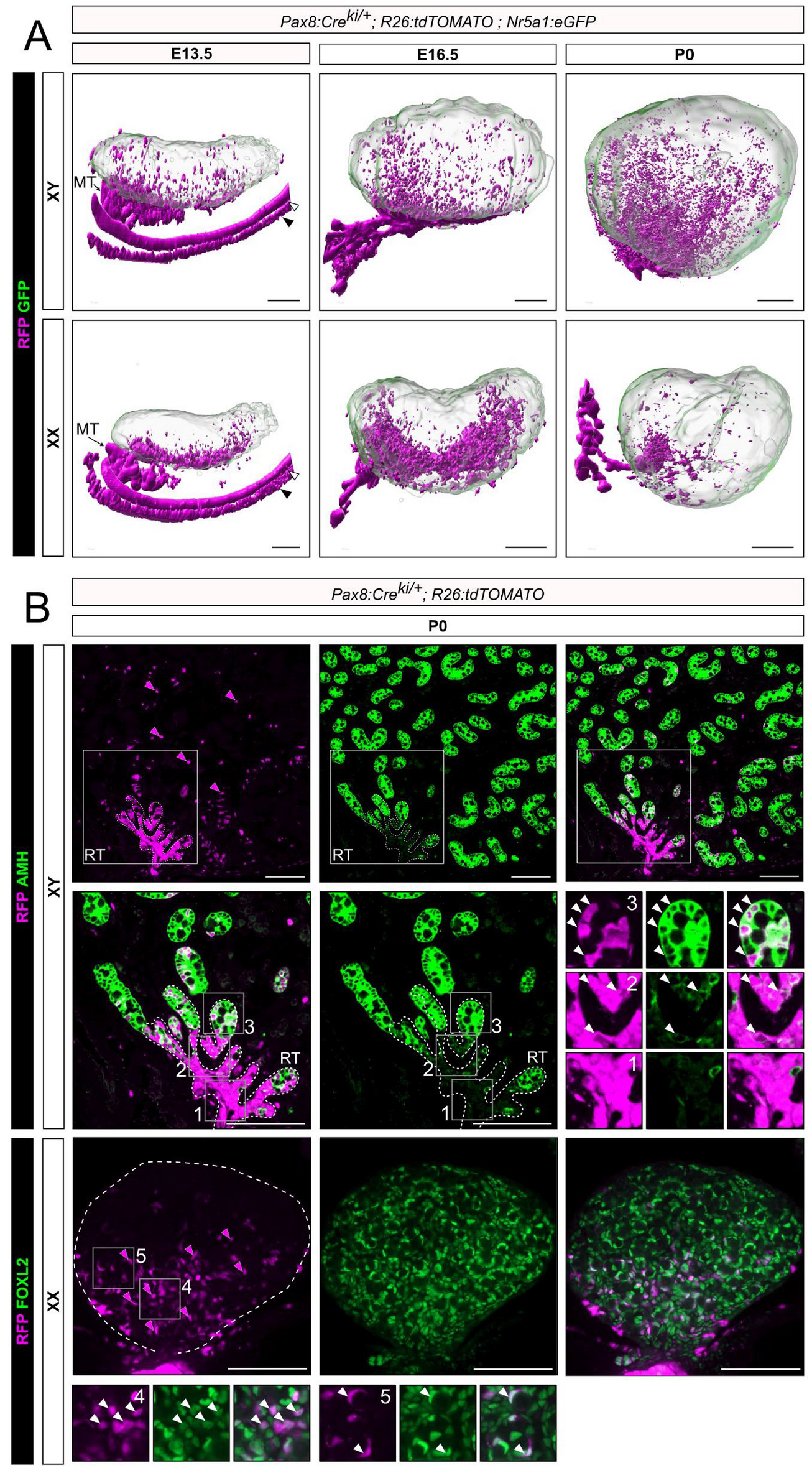
*Pax8*+ progenitors contribute to both rete testis/ovarii and the pool of Sertoli and granulosa cells. (**A**) 3D reconstruction of *Pax8:Cre;Rosa26:tdTomato;Nr5a1:GFP* testis and ovary at E13.5, E16.5 and P0. *Pax8^+^* cells are lineage-traced with RFP. GFP, expressed under the control of the *Nr5a1* promoter, is used to delineate gonadal cells. Note the presence of RFP^+^ cells close to the rete testis/ovarii but also throughout the gonads. White arrowhead indicates the Wolffian duct and empty arrowhead the Müllerian duct. (**B**) Double immunofluorescence for RFP/AMH and RFP/FOXL2 respectively in XY and XX *Pax8:Cre;Rosa26:tdTomato;Nr5a1:GFP* mice at P0. Note the presence of RFP^+^ cells increases near the rete testis and rete ovarii. Cells in the rete testis are exclusively RFP^+^ and do not express AMH (inset 1). At the junction between the rete testis and the testis cords, low AMH expression is observed in some RFP^+^ cells (inset 2). RFP^+^/AMH^+^ cells are also present in testis cords (inset 3). Similarly, RFP^+^/FOXL2^+^ cells are present in granulosa cells of the developing ovary (inset 4) including in some primordial follicles (inset 5). Red arrows indicate RFP^+^ cells, yellow arrows indicate RFP^+^/AMH^+^ or RFP^+^/FOXL2^+^ cells. DAPI was used as a nuclear counterstain. MT, mesonephric tubules; RT, rete testis. Scale bars 200 μm in (**A**) and 100 μm in (**B**).

Lineage tracing experiments using *Pax8:Cre;R26:tdTomato* transgenic animals at E13.5 and E16.5 coupled with IF for SOX9 in the testis and FOXL2 (granulosa cell marker) in the ovary revealed that RFP^+^/SOX9^+^ and RFP^+^/FOXL2^+^ cells are predominantly localized in the cranio-dorsal region of the gonad where the rete develops (**Figure S5**). We also observed some Sertoli and granulosa cells expressing RFP, indicating that *Pax8*-expressing cells contribute also to supporting cells.

To better characterize the ability of *Pax8*-expressing cells to differentiate into rete cells and supporting cells in XX and XY embryos, testes and ovaries of *Pax8:Cre;R26:tdTomato* transgenic animals were analyzed by IF for RFP and AMH (Sertoli cell marker) or FOXL2 (granulosa cell marker) at P0. In the testis, RFP^+^ cells were mainly detected in the rete but a significant number of positive cells were also identified in testis cords (**Figure 5B**). In the rete testis, RFP^+^ cells are mostly AMH^−^ (**Figure 5B** and **inset 1**), while at the interface between the rete testis and the testis cords, some RFP^+^ cells express AMH at a low level (**Figure 5B** and **inset 2**), suggesting that they are Sertoli cells derived from *Pax8*^+^ SLC progenitors. Strikingly, we found that in testis cords, Sertoli cells double positive for RFP and AMH are distributed in a gradient (**Figure 5B** and **inset 3**), with the proportion of doubly positive cells being higher near the rete testis. Double IF analysis revealed that 9.8% of all AMH^+^ Sertoli cells co-express RFP (1,106 double AMH^+^/RFP^+^ cells amongst 11,271 AMH^+^/RFP^−^ and AMH^+^/RFP^+^ cells). Finally, to determine if *Pax8*-expressing cells give rise to other cell types in the developing testis, we evaluated the co-expression of RFP^+^ with HSD3B, a marker of Leydig cells, ACTA2 (also called αSMA), a marker of peritubular myoid cells and vascular smooth muscle cells, ARX, a specific marker of steroidogenic progenitors, and NR2F2 (also called COUP-TFII), a marker of interstitial progenitors. With rare exceptions, none of the RFP^+^ cells co-expressed these markers, consistent with the supporting-like transcriptomic profile of *Pax8*^+^ SLC progenitors (**Figure S6**).

A similar approach was used to assess the capacity of XX SLC progenitors to differentiate into rete ovarii cells and granulosa cells (**Figure 5B**). As expected, RFP^+^ cells were detected mostly in the rete ovarii, but a gradient of positive cells was also identified within the ovary itself. We also found RFP and FOXL2 double-positive cells in a gradient, labeling granulosa cells from ovigerous cords or primordial follicles (**Figure 5B** and **insets 4** and **5**, respectively). Double IF analysis revealed that 6.7% of all FOXL2^+^ granulosa cells co-express RFP (205 double FOXL2^+^/RFP^+^ cells amongst 3,055 FOXL2^+^/RFP^−^ and FOXL2^+^/RFP^+^ cells). RFP^+^ cells co-expressing the granulosa cell marker FOXL2 are also found in the ovary at E13.5 and E16.5 (**Figure S5**). Overall, these results indicate that *Pax8*^+^ progenitor cells are capable not only of forming the rete system but also of giving rise to a significant fraction of the Sertoli and granulosa cell pool.

### The pro-ovary factor WNT4 is a crucial regulator of the SLC lineage and is required for the formation of rete testis

While the signalling pathway(s) responsible for SLC differentiation and rete testis development remain unknown, our scRNA-seq data revealed that in the developing gonads, *Wnt4* is highly expressed in XX and XY SLCs around E11.5, before decreasing (**Figure 6A** and **Figure S3**). To assess whether WNT4 might act in an autocrine manner and activate the WNT/β-catenin signaling pathway in both XX and XY SLCs, we first investigated the temporal expression of *Rspo1* and downstream targets of the WNT/β-Catenin pathway in the developing SLC and supporting cell lineages. As expected, *Rspo1*, *Axin2*, *Sp5*, *Lef1* and *Nkd1* were highly expressed in granulosa cells but almost absent in developing Sertoli cells (**Figure 6A**). By contrast, these genes are expressed in both XX and XY SLCs, indicative of an autocrine effect of WNT4 on SLCs. Initially, at E11.5, the expression levels of *Wnt4* and downstream targets of the WNT/β-catenin pathway are generally similar in both XX and XY SLCs and then a sexual dimorphism appears at E12.5, with higher expression in XX SLCs compared to XY SLCs.

**Figure 6.**
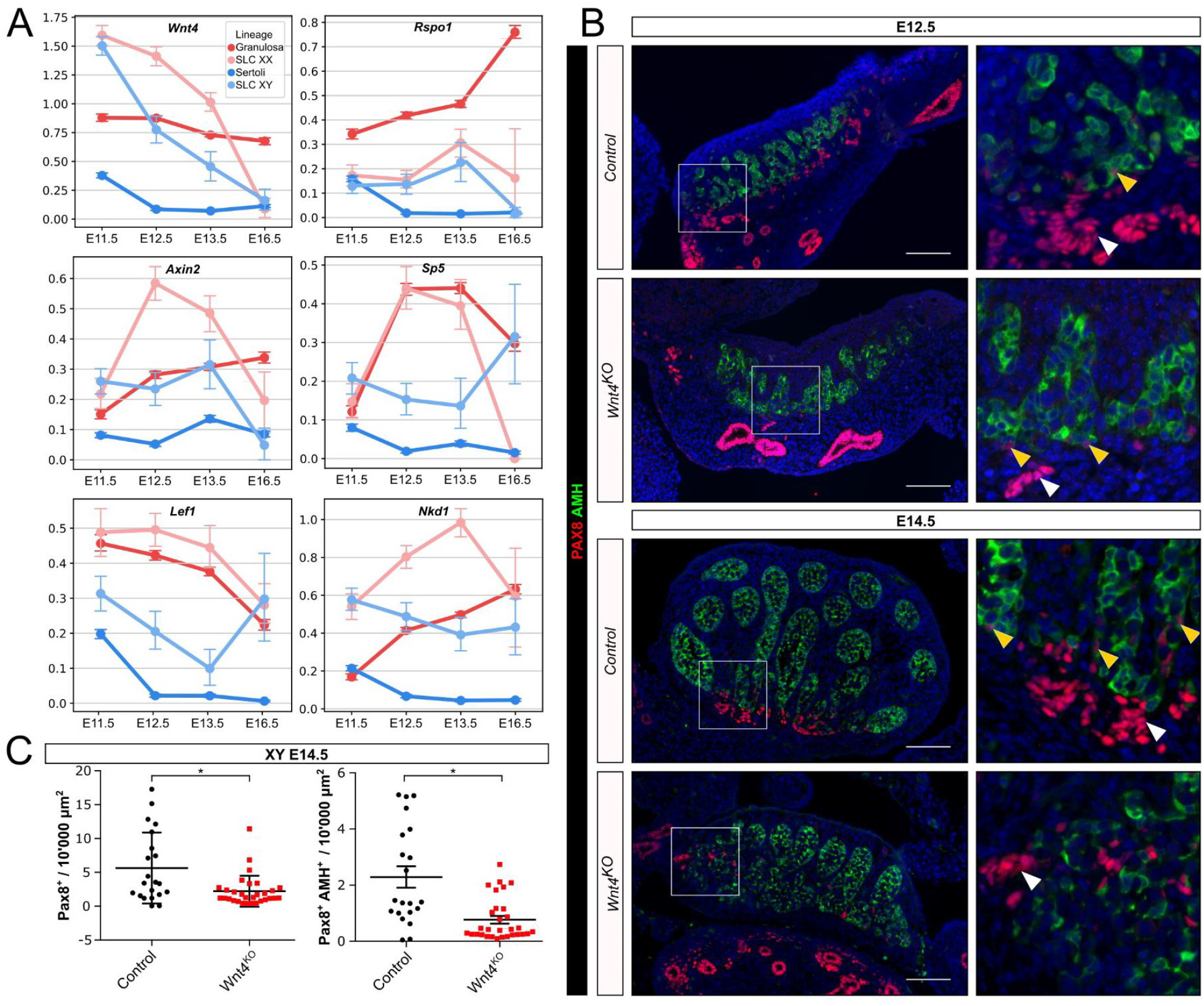
*Wnt4* is required for rete testis formation. (**A**) Expression profiles of genes involved in WNT/β-catenin pathway in SLCs and supporting cells. Data were extracted from scRNA-seq analysis of XX and XY embryos at E11.5, E12.5, E13.5 and E16.5. (**B**) Representative double immunofluorescence against PAX8 and AMH in control (*Sf1:Cre;Sox9^F/+^;Wnt4KO^+/+^*) or *Wnt4*^KO^ (*Sf1:Cre;Sox9^F/+^;Wnt4:KO^−/−^*) XY embryos at E12.5 and E14.5. Boxes indicate regions shown on the right magnifying the rete testis and adjacent testis cords. Note that the number of PAX8^+^ cells, the rete testis and the testis cords near the rete testis are reduced and disorganized in *Wnt4* mutant embryos both at E12.5 and E14.5. White and yellow arrowheads indicate PAX8^+^ and PAX8^+^/AMH^+^ cells, respectively. DAPI was used as a nuclear counterstain. Scale bar 100 μm. (**C**) Quantification of PAX8^+^ and PAX8^+^/AMH^+^ cells per 10,000 μm^2^ of testis section of control and *Wnt4*^KO^ embryos at E14.5. Data are presented as a box-and-whisker plot to illustrate the heterogeneity of PAX8-positive cells according to the different sections. Each point represents the quantification of one testicular section. Three embryos per genotype were used. Student’s t-test, two-sided unpaired (*p < 0.05).

To test the hypothesis that WNT4 is involved in regulating the SLC lineage and rete testis development, we performed double IF for PAX8 and AMH at E12.5 and E14.5 on testes that were either control or homozygous knockout for the *Wnt4* mutant allele (Jeays-Ward et al., 2004; Stark et al., 1994) (**Figure 6B**). As previously reported, we found a reduction in size and fewer testis cords in the *Wnt4* knockout testis (*40*). In addition, we observe a significant reduction in both PAX8^+^ and PAX8^+^/AMH^+^ cells (**Figure 6B** and **C**). The few remaining PAX8^+^ cells are mainly scattered at the base of the testis near the mesonephros but do not form a rete testis. Furthermore, it appears that the testis cords adjacent to the base of the testis and close to the mesonephros are also altered in shape and ill-defined, in particular at E14.5, as if the absence of PAX8^+^ cells and rete testis in *Wnt4* knockout testis prevents the optimal formation of testis cords. Overall, our results reveal that WNT4 is a crucial regulator of the canonical β-catenin signaling pathway in SLCs and is required for the formation of rete testis.

Our scRNA-seq data revealed also that the orphan nuclear receptor DAX1 (encoded by *Nr0b1*) is expressed at high levels in both XX and XY SLCs (**Figure S4**). The function of DAX1 in gonadal development, Sertoli cell differentiation and specification of SLC cells remain unclear and controversial (*21, 41*). DAX1 has been reported to play a role in testis morphogenesis as *Nr0b1*-deficient mice are infertile due to an obstruction of the rete testis and efferent ductules by dysregulated proliferation of Sertoli cells (*42*). In parallel, DAX1 has been identified as an all-trans retinoic acid (ATRA) effector gene. Several observations indicate that ATRA drives *Nr0b1* expression directly, antagonizing the testis determination pathway in somatic lineages (*43–45*). In particular, DAX1 has been reported to antagonize NR5A1 function and impede AMH production in Sertoli cells (*46–48*). We decided to evaluate the hypothesis that abnormalities in ATRA signaling could lead to a defect in the SLC specification and formation of rete testis or rete ovarii. To this end, we investigated the presence of PAX8-positive cells and the structure of the rete testis/ovarii in the gonads of E14.5 embryos ubiquitously lacking all nuclear retinoic acid receptors (RARA, RARB and RARG isotypes) in all cell-types from embryonic day 11.5 onwards (*49*). Our results revealed that RARs, and therefore endogenous ATRA, does not affect SLC numbers and are dispensable for rete testis and rete ovarii formation (see **Figure S7**). Nevertheless, it still remains possible that DAX1 may have a role in mediating SLC lineage specification and differentiation.

## Discussion

Although the process of gonadal sex determination has been studied for several decades, many key aspects remain elusive. Our understanding of gonad development and differentiation will only be complete after full characterization of all its cell types. We are still missing a comprehensive understanding of the different cell lineages comprising this bipotential primordium over time and their relationships to each other. Similarly, molecular factors governing the specification of the many cell lineages that emerge from this process are still missing. The widespread adoption of advanced sequencing technologies such as scRNA-seq, has provided the field of developmental biology with an opportunity to discover previously unrecognized cell types, such as short-lived progenitors or rare cell lineages.

Here, we densely sampled gene expression at the single-cell level in developing mouse gonads during the critical period of sex determination. We describe the specification and differentiation of a rare, previously overlooked, gonadal cell lineage. We refer to these as supporting-like cells (SLCs) due to their transcriptomic similarities to the supporting cell lineage. Transcriptionally early SLCs are closely related to CE-derived progenitors and are specified at E10.5, prior to sex determination. These cells are localized in the XX and XY genital ridge along the border to the mesonephros and express *Pax8* as well as CE-derived markers, such as *Nr5a1*, *Gata4* and *Wt1*. We found that SLCs are initially sexually undifferentiated and co-express both the pro-ovarian and pro-testis genes *Wnt4* and *Sox9*. From E12.5 onward sexual dimorphism appears with the gradual acquisition of a Sertoli-like and granulosa-like profiles. Finally, our lineage tracing experiments revealed that *Pax8*^+^ progenitors contribute primarily to the formation of the rete testis and ovarii and also to a significant fraction of the Sertoli and granulosa cell pool.

### Origin and specification of SLCs

Our transcriptomic analyses reveal that SLCs are the first somatic cells of the gonad to be specified, as early as E10.5. By contrast, the commitment of CE-derived progenitors toward the supporting lineage is initiated just before the process of gonadal sex determination at E11.0 - E11.5 (*7*). We found that at E10.5, early somatic progenitor cells - known to express marker genes such as *Gata4*, *Nr5a1* and *Wt1* - are not a homogeneous population. Using scRNA-seq, we identified several CE-related sub-populations with close but distinct transcriptomic profiles, including one that expresses the SLC markers *Pax8, Ncam1* and *Enpp2*, indicating already a transition toward early and late SLCs. At this early stage (E10.5), the transcriptomes of *Pax8*^−^ and *Pax8*^+^ somatic progenitor sub-populations are extremely similar with only 112 differentially expressed genes including the SLC markers *Pax8*, *Ncam1* and *Enpp2* out of a total of about 15,000 genes (see **Data S3**). This single cell analysis strongly suggests that the SLC lineage originates and diverges transcriptionally as early as E10.5 from somatic progenitor cells expressing the CE marker genes *Gata4*, *Nr5a1* and *Wt1*. These findings are consistent with our IF results, revealing that at E11.0, a population of PAX8^+^ cells co-expressing GATA4 is localized at the base of the genital ridge in close contact with the mesonephric tubules (see **Figure 4, Figure S2**). While these PAX8^+^ cells are initially distributed all along the gonad at the border to the mesonephros, they are gradually restricted to the anterior part in close contact with the cranial mesonephric tubules. Our results are also in agreement with the previous description of a population of cells expressing NR5A1 found in both sexes between the cranial mesonephric tubules and the undifferentiated gonad prior to sex determination (*22*). Finally, the transcriptomic similarities between SLCs and the supporting lineage, as well as the expression of Sertoli (*Dhh, Ptgds, Aard, Nedd9, Tesc*) and granulosa (*Foxl2, Irx3, Fst*) markers by rete testis and rete ovarii cells respectively, reinforce the hypothesis of a CE origin for the SLC lineage. Collectively, our data indicate that SLCs are derived from the CE and share a common origin with the supporting cell lineage.

### *Pax8*^+^ progenitors contribute to both rete testis/ovarii and to the pool of Sertoli and granulosa cells

By coupling lineage tracing analysis with immunofluorescence, we observed three distinct RFP^+^ cell types in the developing testis. The first type is RFP^+^/AMH^−^ and is localized in the center of the rete testis (**Figure 5G-K** and **inset 1**). These cells form the core network of interconnected tubules of the rete testis. The second population expresses RFP and low levels of AMH and is located at the interface between the rete testis and the testicular cords (**Figure 5G-K** and **inset 2**). These cells most certainly correspond to the transitional zone and/or the tubuli recti, both of which are composed exclusively of Sertoli cells (*20, 50, 51*). The third population is composed of Sertoli cells that are located within the testis cords and represent 10% of all Sertoli cells at birth (**Figure 5G-K** and **inset 3**). These results indicate that *Pax8*^−^expressing cells contribute not only to the rete testis but also significantly to the Sertoli cell pool, suggesting a potential unappreciated role for PAX8 in supporting progenitors. However, the remaining question is whether the RFP^+^ Sertoli cells are derived solely from the 5% pre-supporting progenitors that weakly express *Pax8* around E11.5 (see **Figure 2I**), or whether, remarkably, they are derived in part or exclusively from the SLC lineage. Our lineage tracing experiment does not allow us to give a definitive answer to this question. Nevertheless, several indications suggest the possibility that SLC progenitors could be a second source of Sertoli cells. First, both the SLC and supporting lineages appear to derive from a common progenitor at around E10.5, which expresses CE-derived markers (see **Figure 3A, C** and **D**). Furthermore, the transcriptome of SLCs shows similarities with cells of the supporting lineage, both at early stages with pre-supporting cells and at later stages with granulosa and Sertoli cells (see **Figure 2I** and **J**). For example, from E12.5 onwards, SLCs acquire either a Sertoli- or granulosa-like identity with overexpression of many classical Sertoli or granulosa markers (**Figure S4)**. Furthermore, our lineage tracing analysis of *Pax8*^+^ cells reveals a gradient of RFP^+^ Sertoli cells in the testis, with a higher density near the rete testis, suggesting that SLC progenitors may indeed contribute to these RFP^+^ Sertoli cells (**Figure 5A-B**). The ability of these potential SLC-derived Sertoli cells to colonize all regions of the testis reflects their early inclusion in the expanding testis cords following the rapid proliferation of fetal Sertoli cells. Finally, it also remains possible that at around E10.5 the specification of some early SLC progenitors is not yet well established redirecting a small fraction of these cells to a supporting fate instead. This could explain the transient presence of *Pax8* transcripts in 5% pre-supporting progenitors around E11.5.

In the developing ovary, our results revealed that RFP^+^ cells have the ability to differentiate into either rete ovarii cells or granulosa cells. The RFP^+^/FOXL2^−^ cells forming the rete ovarii are concentrated mostly in the cranio-dorsal part of the ovary consistent with the IF and light sheet microscopy at E13.5 and E16.5 (**Figure 5A-B** and insets **4** & **5**). Regarding granulosa cells, our quantitative analyses reveal that about 6% of FOXL2-expressing granulosa cells are derived from *Pax8*^+^ progenitors. As for the testis, we observe a gradient of these cells, with RFP^+^/FOXL2^+^ double-positive cells mainly concentrated in the cranio-dorsal part of the ovary near the rete ovarii. In the cortical part, we also observe primordial follicles with some granulosa cells being RFP^+^.

### Factors regulating SLC specification, differentiation and rete formation

Although our combined scRNA-seq analysis and lineage tracing approach has allowed us to better characterize the origins and developmental trajectories of the SLC progenitor cells that give rise to the rete system, some questions still remain. In particular, the factors, local signals and molecular pathways responsible for the specification of SLC progenitors remain unknown. We decided to focus our attention on the secreted protein WNT4 because of its high expression in the SLC lineage (**Figure S3**) and the essential role of the canonical WNT/β-catenin signaling pathway for ovarian fate, particularly in the supporting cell lineage (*52–56*). We tested the possibility that WNT signaling also plays a role in the SLC lineage and the formation of the rete testis. Indeed, the high expression of *Wnt4* and its downstream target genes (*Lef1*, *Axin2*, *Sp5* and *Nkd1*) in the SLC lineage is consistent with an autocrine effect (**Figure 6A**). Direct experimental evidence found in the literature already suggested that canonical β-catenin signaling is activated in the rete ovarii/testis of the gonads at E12.5 (*52*). Using a reporter line carrying *LacZ* fused to *Axin2* (*57*), a target of β-catenin, Chassot and coworkers observed that at E12.5, a robust β-galactosidase staining was found at the interface between the XX and XY gonads and the mesonephros, a position corresponding to the rete testis and rete ovarii at this developmental stage.

Our results in *Wnt4* knockout mutant mice show a drastic reduction in the number of SLCs at E14.5, as well as disorganization of the rete testis and adjacent testicular cords (**Figure 6B**). This confirms that WNT4 and the canonical WNT/β-catenin signaling pathway are required for rete testis formation. However, it is not clear whether the reduction in the number of PAX8^+^ cells and the alteration of the rete testis formation is due to a reduction in the total number of SLCs or to a change in their fate. The former is unlikely, as our transcriptomic data combined with results from EdU DNA synthesis monitoring assay clearly indicate that SLCs do not proliferate during the process of testicular determination. This absence of proliferation may explain why the rete system is a relatively small structure compared to the whole gonad (**Figures 2J** and **S2**). We hypothesize instead that the ability of progenitors to maintain their SLC identity and form the rete testis is dependent on the canonical WNT/β-catenin pathway. Loss of the WNT/β-catenin pathway may therefore affect the fate of SLC progenitors. Since the rete testis forms an essential bridging system connecting the testis cords to rete testis, an indirect effect is the alteration of the morphology of the testicular cords in close proximity to the defective rete structure, as observed in *Wnt4* knockout mutant testes. If the WNT pathway is indeed key to the differentiation of SLC progenitors towards rete testis fate, what is their fate in its absence or upon overexpression of *Wnt4* or the canonical WNT pathway more generally? An increase in the WNT/β-catenin pathway could be tested by genetic studies promoting stabilization of β-catenin or ablation of *Znrf3*, a negative regulator of ovarian development. Loss of *Znrf3* results in ectopic canonical WNT signaling in XY gonads with reduced *Sox9* expression, resulting in defects in testis determination, including gonadal sex reversal (*58*). We predict that elevated WNT/β-catenin signaling would promote SLC fate and increase the final number of *Pax8*^+^ SLCs in the XY mutant ovary/ovotestis.

Overall, our results provide a single-cell transcriptional atlas of gonadal sex determination in mice. This unique dataset allowed us to identify SLCs as a previously uncharacterized cell population at the origin of the rete system. Understanding this lineage is important for two reasons: first, proper formation of the rete is essential for male fertility, as it provides a channel system allowing sperm export from the testis to the efferent ductules. Second, from a fundamental point of view, the SLC lineage exhibits remarkable characteristics, and represents an interesting model for the study of cell fate decisions during mammalian organ development. We demonstrate that the SLC lineage is the first somatic cell lineage to be specified, even before the initiation of the gonadal sex determination process. We are unable to definitely demonstrate that the SLC lineage possesses the capacity to differentiate into Sertoli-like or granulosa-like cells, a feature that was previously thought to be unique to the supporting cell lineage. However, several observations support this conclusion. The molecular mechanism involved in Sertoli-like differentiation is different from that of the supporting lineage, since it does not involve the well-known pro-testis gene *Sry*. It is now essential to identify the factors that control the specification of the SLC lineage as well as the molecular mechanisms controlling their differentiation into rete cells exhibiting transcriptomic profiles close to Sertoli or granulosa cells. In addition, a better characterization of the similarities and differences between the supporting and SLCs lineages during the process of testis/ovarian development is needed. With our results, we lay the foundation for further elucidation of mechanisms in mammalian sex determination, and improving our understanding of the genetic basis and physiology of human disorders/differences of sex development.

## Material & Methods

### Animals

Animals were housed and cared according to the ethical guidelines of the Service de la Consommation et des Affaires Vétérinaires (SCAV) of the Canton de Genève (experimentation ID GE214-19 and GE35) or the relevant institutional and European animal welfare laws, guidelines and policies for the *Wnt4*^KO^ line. *Pax8*^*tm1.1(cre)Mbu*^/*J:Gt(ROSA)26Sor*^*tm14(CAG-tdTomato)Hze*^:*Tg(Nr5a1/EGFP)1Klp* (abbreviated *Pax8*:*cre*;*Rosa26*:*Tomato*;*Nr5a1*:*eGFP*) mouse strains were described previously (*30, 39, 59*). These mice have been maintained on a mixed 129/CD1 genetic background and were genotyped between P6 and P11 from digit biopsies by PCR as described previously (*39*). For *Pax8:cre* knock-in allele, primers used were 5’-TCTCCACTCCAACATGTCTGC-3’, 5’-AGCTGGCCCAAATGTTGCTGG-3’ (*30*) and 5’-GCCAGCAGCTATGAGGTTGA-3’, giving a WT band of 601 bp and a specific Cre amplicon of 673 bp. Control (*Sf1:Cre;Sox9^F/+^;Wnt4KO^+/+^*) and *Wnt4*^KO^ mutant (*Sf1:Cre;Sox9^F/+^;Wnt4:KO^−/−^*) embryos were generated and genotyped as described in (*60*) and the mouse line was kept on a mixed 129/C57Bl6/J background.

### Sample collection and single cell transcript profiling

The day when a vaginal plug was designated as embryonic day E0.5. Embryos were collected at E10.5 (8±2 caudal somites), E11.5 (19±4 ts), E12.5, E13.5, and E16.5 from time-mated pregnant female CD-1 outbred mice (Charles River) and heterozygous Tg(Nr5a1-GFP) transgenic male mice (*39*). The *Nr5a1-GFP* transgene was used to facilitate identification of the position of the genital ridges during the dissection. Genetic sexing was performed by PCR, as described previously (*61*). Urogenital ridges including both the gonads and mesonephros (E10.5, E11.5) from both sexes, and testes or ovaries (E12.5-E16.5), were isolated at each time point and enzymatically dissociated as described previously (*62*). Approximately 5000 single cells were loaded on a 10x Chromium instrument and scRNA-seq libraries were prepared using the 10X Genomics Chromium Controller and Chromium Single Cell 3’ Library & Gel Bead Kit V2, as per the manufacturer’s instructions. For each developmental stage and sex, two independent biological replicates were processed. The prepared libraries were sequenced on an Illumina HiSeq4000 using paired-end 26 + 98 + 8 bp as the sequencing mode. Libraries were sequenced at a targeted depth of 100 000 to 150 000 total reads per cell. Sequencing was performed at the Health 2030 Genome Center, Geneva.

### Data pre-processing with the Cell Ranger package, cell selection and quality controls

Computations were performed at the Vital-IT Center for high-performance computing of the SIB (Swiss Institute of Bioinformatics) (http://www.vital-it.ch). Demultiplexing, alignment, barcode filtering and UMI counting were performed with the Cell Ranger v2.1 pipeline (10x Genomics). Data were mapped to the mouse reference genome GRCm38.p5 in which the eGFP (NC_011521.1) combined with the bovine GH 3’-splice/polyadenylation signals (*39*) (NM_180996.1) sequences have been added. It has recently been established that the mouse *Sry* locus harbors a cryptic exon that is essential for male sex determination (*63*). As this exon is located in a palindromic region, all reads mapping to this region are also mapping to the positive strand and discarded as multimappers. In order to include reads from exon 2 of the *Sry* gene in our counts matrices, reads from the negative strand, mapped to location chrY(2653159, 2655636) with number of alignments (NH) tag inferior to 3 were extracted and GN tag was set to “Sry”. The edited reads were saved in a specific bam and reads were deduplicated and counted using umi-tools count function with --extract-umi-method=tag --per-cell --cell-tag=CB --per-gene --gene-tag=GN –umi-tag=UB parameters.

To select barcodes associated to cells, we set a threshold as the local minima of a standardized density curve of UMI counts located between the knee point and the inflection point of the ranked barcodes distribution plot (DropletUtils package). When no local minimum could be detected between the two points, the nearest local minimum was used. Quality controls regarding abnormal mitochondrial or ribosomal content, UMI number, detected gene numbers, unmapped reads and putative doublet identification (Scrublet 0.2.1-0) were performed, but no data filtering was applied as no important anomalies were detected. In total, for gonadal samples, we obtained 92,267 cells. It included 13,060 cells from E9.0, 14,962 cells from E10.5, 16,670 cells from E11.5, 20,285 cells from E12.5, 25,794 cells from E13.5 and 16,994 cells from E16.5. In total, for gonadal samples, we obtained 94,705 cells. It included 14,962 cells from E10.5, 16,670 cells from E11.5, 20,285 cells from E12.5, 25,794 cells from E13.5 and 16,994 cells from E16.5. In total, we obtained 53,962 XY cells and 40,743 XX cells.

### Gene expression normalization

UMI counts per gene per cell were divided by the total UMI detected in the cell with exclusion of very highly expressed genes (accounting to more than 5% of total counts in cell), multiplied by a scale factor of 10,000 and log transformed.

### Dimensionality reduction, Batch correction, Clustering and cell-type annotation

A first step of dimensionality reduction was performed using Independent Component Analysis (icafast function, ica package version 1.0-2) on all genes expressed in more than 50 cells (100 components). Because ICA extracts non-Gaussian components from data and creates non-orthogonal components (as in PCA) it allows a better discrimination of small cell populations in heterogeneous dataset. We set the number of ICs to 100 as we expected to have fewer than 100 different populations in our total dataset. Another advantage of ICA is its good performance with nearly no gene filtering. We did not perform any highly variable gene selection.

To minimize biological or technical variability on clustering, we created a neighbor graph based on ICA components using the BBKNN python module (version 1.3.12) (*64*). The correction was performed between biological replicates. BBKNN corrects the neighbor graph by identifying the k-nearest neighbors in each batch instead of considering all cells come from one unique batch. No correction of the expression levels was performed.

A first clustering was performed using the Scanpy Leiden method with resolution 1.3. Clusters were annotated using known markers from described cell populations of developing testis and ovary. To refine the clustering, a leiden clustering was applied to the SLC cluster and granulosa cells to obtain early and late SLC. The parameters to achieve this final clustering of the cells were resolution=0.37, restrict_to=(’leiden’, [ “5”, “30”]).

### Lineage reconstruction, UMAP visualization

Lineage reconstruction was done using partition-based graph abstraction (PAGA) (*65*). From the global k-nearest-neighbors graph and clustering, PAGA outputs a much simpler graph with nodes representing clusters and edges showing confidence of connection between clusters. PAGA and its visualization (Fruchterman-Reingold) were run prior to UMAP to allow a better organization of clusters. For complete dataset global visualization, UMAP was run with random_state=25, min_dist=0.2, init_pos =“paga” parameters. (**Figure 1**, **Figure S1**).

In order to precisely reconstruct the SLC cell lineage, we created three subsets of data: the first two groups contain XX or XY cells from clusters composed of cells from the coelomic epithelium, surface epithelium, pre-supporting cell population, granulosa and Sertoli cells as well as early and late SLC. The third subset is composed of XX and XY cells from clusters composed of cells of the coelomic epithelium, as well as early and late SLCs. Clusters with less than 50 cells were removed ( a few XY cells clustering with granulosa cells for instance). We observed that in some conditions PAGA outputs stronger connections with small clusters. To address this issue, we randomly subsampled big clusters to obtain balanced clusters and ran PAGA on this subsampled data. BBKNN, PAGA and UMAP steps were re-ran independently on XX, XY and SLC datasets.

### Correlation analysis

The similarity of the transcriptomes between the different cell populations was determined using Spearman correlation analysis on the mean expression of the genes. We then applied a hierarchical clustering on the correlation matrix using ward linkage (*66*). The obtained dendrogram was reused to order cell types in figure 1J.

### Differential expression analysis

Differential expression analysis was performed using the Seurat (version 3.2.2) wrapper function for MAST (version 1.14.0). Genes were selected with a log-foldchange higher than 0.25 and an adjusted *p*-value below 0.05. Gene ontology enrichment was performed positive or negative genes obtained with differential expression analysis using clusterProfiler package (version 3.16.1) and only on Gene Ontology biological processes database. Background was set to genes detected in more than 50 cells in the complete dataset and *p*-value cutoff was set to 0.01.

### Scoring of the cells

To create a classification score for the different lineages of gonad specific cells, we trained a Least Absolute Shrinkage and Selection Operator (LASSO) with cross-validation classifier on a random subset of cells with a one versus all approach. The LASSO approach uses a L1 regularization and allows a stringent selection of the genes defining the cells of interest (https://www.jstor.org/stable/2346178). We used the LassoCV function from scikit-learn python module. To avoid gene weighting bias due to over-representation of some clusters, the cells were randomly subsampled keeping 300 cells per leiden clusters.

### Histological and Immunofluorescence analyses

Embryonic or postnatal samples were collected at specific time points determined by the day of vaginal plug (E0.5) and fixed overnight at 4°C in 4% paraformaldehyde. Samples were embedded in paraffin and 5μm sections were prepared. Unmasking was performed in TEG (10mM, pH 9) or citrate (10 mM, pH 6) buffer for 15 minutes in a pressure cooker. Sections were incubated in blocking buffer (3% BSA, 0.1% Tween in PBS) for 2 hours at room temperature. Primary antibodies were incubated overnight at 4°C. Secondary antibodies were incubated 1 hour at room temperature with DAPI (1:1000). The following primary antibodies were used: sheep anti-SOX9-CT (1:200, a generous gift from Prof. Francis Poulat, University of Montpellier, France), rabbit anti-FOXL2 (1:300, (*67*)), mouse anti-αSMA (1:500, a generous gift from Christine Chaponnier, University of Geneva, Switzerland), rabbit anti-ARX (1:500, a generous gift from Prof. Ken-ichirou Morohashi, Kyushu University, Japan), goat anti-AMH (C-20) (1:100, sc-6886, SantaCruz Biotechnology, USA), mouse anti-AMH (1:20, MCA2246, Bio-Rad Laboratories), rabbit anti-PAX8 (1:200, 10336-1-AP, Proteintech), mouse anti-RFP (DsRed) (1:250, sc-390909, SantaCruz Biotechnology, USA), goat anti-RFP (1:50, 200-101-379, Rockland Immunochemicals Inc., USA), rabbit anti-RFP (1:250, 600-401-379, Rockland Immunochemicals Inc., USA), mouse anti-COUPTFII (1:200, PP-H7147-00, Perseus Proteomics Inc., Japan), rabbit anti-HSD3B (1:200, KO607, TransGenic Inc, Japan). Each immunostaining run included negative controls, with replacement of the primary antibody by blocking solution. Fluorescent images were acquired using an Axio Imager M2 or Z1 microscope (ZEISS, Germany) fitted with an Axiocam 702 mono camera or MRm camera (ZEISS, Germany). Images were minimally processed for global levels with ZEN (ZEISS, Germany). Whole gonad images were analyzed using QuPath v0.2.0 (*68*). The percentages of RFP^+^/AMH^+^ or RFP^+^/FOXL2^+^ co-labeled cells were determined from three independent animals. With respect to *Wnt4*^KO^ embryos, paraffin sections covering the entire gonad were processed for immunostaining and the number of PAX8^+^ cells or PAX8^+^ AMH^+^ cells was determined every 20μm using the cell counter Plugin from Fiji. The graphs show the number of positive cells per 10 000μm2 of gonadal area measured in each section, the mean and s.e.m for control (n=3) and *Wnt4*^KO^ mutant embryos (n=3).

### Whole-mount immunofluorescence

For whole-mount immunofluorescence (IF), embryonic samples were fixed overnight at 4°C, rinsed three times for 20 minutes with PBS 0.1% Triton X-100 (PBST) and serially dehydrated into 100% methanol for storage at −20°C. Immunofluorescence was performed as follows: samples were rehydrated stepwise into PBS, washed three times with PBST, rocked for 1 hour at room temperature in blocking solution (BS: 10% FBS, 3% BSA, 0.2% Triton X-100 in PBS), and incubated with primary antibodies in BS overnight at 4°C. The following day, samples were washed three times for 30 minutes in washing solution (WS: 1% FBS, 3% BSA, 0.2% Triton X 100 in PBS) and incubated with secondary antibodies and DAPI (1:1000) in BS overnight at 4°C. The next day, samples were washed three times in WS and mounted in DABCO (Sigma Aldrich, St Louis, USA). Images were taken with an LSM710 confocal microscope using the associated Zen software (Zeiss). The following primary antibodies were used: rabbit anti-PAX8 (1:500, 10336-1-AP, Proteintech), mouse anti-GATA4 (1:500, Sc-25310, Santa Cruz Biotechnology), goat anti-SOX9 (1:1000, AF3075, R&D Systems), rat anti-LAMB1 (1:500, RT-796-P, NeoMarkers), mouse anti-NR2F2 (1:250, PP-H7147-00, Perseus Proteomics).

### EdU labeling

For cell proliferation experiments, pregnant females were injected intraperitoneally with 25 mg/kg 5-ethynyl-2′-deoxyuridine (EdU; Lumiprobe) dissolved in PBS. Embryonic samples were collected 1 hour after EdU injection and fixed overnight at 4°C. EdU treated samples were imaged using the wholemount immunofluorescence protocol with the addition of a click reaction step between rehydration and blocking. Samples were incubated in click reaction solution (20 mg/mL ascorbic acid, 2mM cupric sulfate, 4 μM sulfo-Cy3 azide dye (Lumiprobe) in PBS) for 1 hour rocking at room temperature.

### Sample clearing and light sheet fluorescence microscopy (LSFM)

*Pax8:Cre;Rosa26:tdTomato;Nr5a1:eGFP* embryos at E11.5, E12.5, E13.5, E16.5 were cleared using a modified passive CLARITY-based clearing protocol (*69*) while the samples at embryonic stages E10.5 and dissected gonads at P0 went through a refractive-index matching process. Briefly, samples at E11.5, E12.5, E13.5 and E16.5 embryonic stage were incubated in 5 mL X-CLARITY™ Hydrogel Solution Kit (C1310X, Logos Biosystems, South Korea) for 3 days at 4°C, allowing diffusion of the hydrogel solution into the tissue. Polymerization of solution was carried in a Logos Polymerization system (C20001, Logos Biosystems, South Korea) at 37°C for 3 hours. After two washes of 30 min in PBS, samples were immersed in a SDS based clearing solution and left at 37°C for 24h (E11.5), 48h (E12.5), 72h (E13.5) and 96h (E16.5). Once cleared, tissue was washed twice in PBS-TritonX 0.1% and then placed in a Histodenz© based-refractive index-matching solution (RI = 1.46). Early-stage embryos (E10.5) underwent a uniquely refractive index-matching process, prior to 2 days of incubation in PBS-TritonX 2% at room temperature (RT). Dissected gonads of embryos at P0 were embedded in low melting agarose, placed in PBS-TritonX 2% for 2 days at RT and transferred in a Histodenz© based-refractive index-matching solution (RI = 1.46) 2 days before imaging. Light-sheet imaging was performed using a customized version of the Clarity Optimized Light-sheet Microscope (COLM) (Tomer et al., 2014) at the Wyss Center Advanced Light-sheet Imaging Center, Geneva. The samples were illuminated by one of the two digitally scanned light sheets, using a 488 nm and 561 nm wavelength laser. Emitted fluorescence was collected by 10X XLFLUOR4X N.A. 0.6 filtered (525/50 nm and 609/54 nm Semrock BrightLine HC) and imaged on an Orca-Flash 4.0 LT digital CMOS camera at 4 fps, in rolling shutter mode. A self-adaptive positioning system of the light sheets across z-stacks acquisition ensured optimal image quality over the whole thickness of the sample. Z-stacks were acquired at 3 μm spacing with a zoom set at 10x resulting in an in-plane pixel size of 0,59 μm (2048 x 2048 pixels). Images of multiple tiles were reconstructed in 3D using the Grid Collection Stitching plugin tool in TeraStitcher (BMC Bioinformatics, Italy). Volume reconstructions and 3D renderings were performed using Imaris Bitplane software.

### Statistical analyses

The number of PAX8^+^ cells or PAX8^+^AMH^+^ cells per unit of gonadal area in control and Wnt4^KO^ mutant embryos was analyzed by a two-tailed t-test using Graph Pad Prism version 9.

## Acknowledgments

We thank Deborah Penet for the sequencing, Christelle Borel (GEDEV department, University of Geneva) for her advice and help with 10X technology, the team of the Animal Facility and the Histology Plateform (Faculty of Medicine, University of Geneva). We thank also the members of the Nef laboratory for helpful discussion and critical reading of the manuscript.

## Funding

Swiss National Science Foundation grant 31003A_173070 (SN)

Swiss National Science Foundation grant 310030_200316 (SN)

Département de l’Instruction Publique of the State of Geneva (SN)

Agence Nationale de la Recherche grant ANR-19-CE14-0022-01 SexDiff (MCC)

UK Medical Research Council grant MC_U142684167 (AG)

National Institutes of Health grant 1R01HD090050 (BC, CB)

National Institutes of Health grant R37HD039963 (BC)

NBG and CD were supported by grants from Agence Nationale de la Recherche (ANR-16-CE14-0017 and ANR-20-CE14-0022)

## Author Contributions

SN, DW and AG contributed to conception and design of the study. IS, YN, and FK collected mouse samples and prepared sequencing libraries. CM, PS, RR, SG, MS performed the bioinformatics analysis. VR, APG, CB, PS, DC, PS, CD, IG, FT and FK carried out mouse work, cell lineage tracing, immunofluorescence, light sheet microscopy and other experimental analysis. CM, VR and SN wrote the first draft of the manuscript. All authors contributed to manuscript revision, read, and approved the submitted version. Funding and Resources, SN, AG, BC, MCC, LB, NBG and DW; Supervision, SN.

## Data and materials availability

All data needed to evaluate the paper are present in the paper and/or the Supplementary Materials. Additional data related to this paper may be requested from the authors.

