## Supplementary Material for "Origin, specification and differentiation of a rare supporting-like lineage in the developing mouse gonad"

**This PDF file includes:**

Supplementary Material& Methods

Figs. S1 to S7 and legends

Table S1

Legends for data S1 to S4

References

**Other Supplementary Material for this manuscript includes the following:**

Data S1 to S4

**Supplementary Material & Methods**

### Whole-mount in situ hybridization

Whole-mount in situ hybridization (WISH) analysis of embryonic gonads and probes for Sox9 and Wnt4 have been previously described (Bogani, et al., 2009, Warr, et al., 2009). At least three independent biological samples from XX and XY embryos were analyzed for each gene.

Inactivation of Rar coding genes

To inactivate *Rar* coding genes, mice bearing *lox*P-flanked (L2) alleles of *Rara* and *Rarg* and null (L-) alleles of *Rarb* were crossed with mice bearing the ubiquitously expressed, tamoxifen-inducible, Cre/ERT^2^ recombinase. Noon of the day of a vaginal plug was taken as 0.5-day embryonic development (E0.5). To activate Cre/ERT^2^ in embryos, one dose of tamoxifen (130 mg/kg body weight) was administered to the pregnant females at E10.5 by oral gavage. Following collection at E14.5, the tail of each embryo was sampled and used for genomic DNA extraction and genotyping as described (Vernet, et al., 2020). The embryos were then fixed overnight in cold 4% (w/v) paraformaldehyde (PFA) solution made in phosphate buffered saline (PBS). They were rinsed in PBS, placed in 70% (v/v) ethanol, and then embedded in paraffin. Consecutive, sagittal, 5µm-thick sections were cut throughout entire gonads. For FOXL2, PAX8 and SOX9 detection, demasking was performed for 1 hour at 95°C in 10 mM sodium citrate buffer at pH 6.0. Sections were rinsed in PBS, and then incubated for 16 hours at 4°C in a humidified chamber with the appropriate primary antibodies (see Table below) diluted in PBS containing 0.1% (v/v) Tween 20 (PBST). After rinsing in PBST (3 times for 3 minutes each), sections were incubated for 45 minutes at 20°C in a humidified chamber with Cy3-conjugated or Alexa Fluor 488-conjugated secondary antibodies (see Table below). Nuclei were counterstained with 4',6-diamidino-2-phenylindole (DAPI) diluted at 10 μg/ml in the mounting medium (Vectashield, Vector Laboratories). For RARG detection, ImmPRESS® Polymer Detection Kit MP-7800-15 was used according to the manufacturer’s protocol (Vector Laboratories). *Rara* and *Rarb* ablations were assessed by PCR analysis of genomic DNA extracted from 5 sections made at the level of the gonad and obtained from each embryo.

| **Antigen** | **Species** | **Reference** | **Source** | **Dilution** | **Secondary antibody or Detection** |
| --- | --- | --- | --- | --- | --- |
| FOXL2 | Goat | ab5096 | Abcam | 1/200 | Alexa Fluor 488-conjugated donkey anti-goat |
| SOX9 | Rabbit | AB5535 | Millipore | 1/1000 | Alexa Fluor 488-conjugated donkey anti-rabbit |
| PAX8 | Mouse | ab53490 | Abcam | 1/10 | Cy3-conjugated donkey anti-mouse |
| RARG | Rabbit | 8965S | Cell Signaling | 1/200 | ImmPRESS® Polymer Detection Kit MP-7800-15 |

Supplementary Figure S1

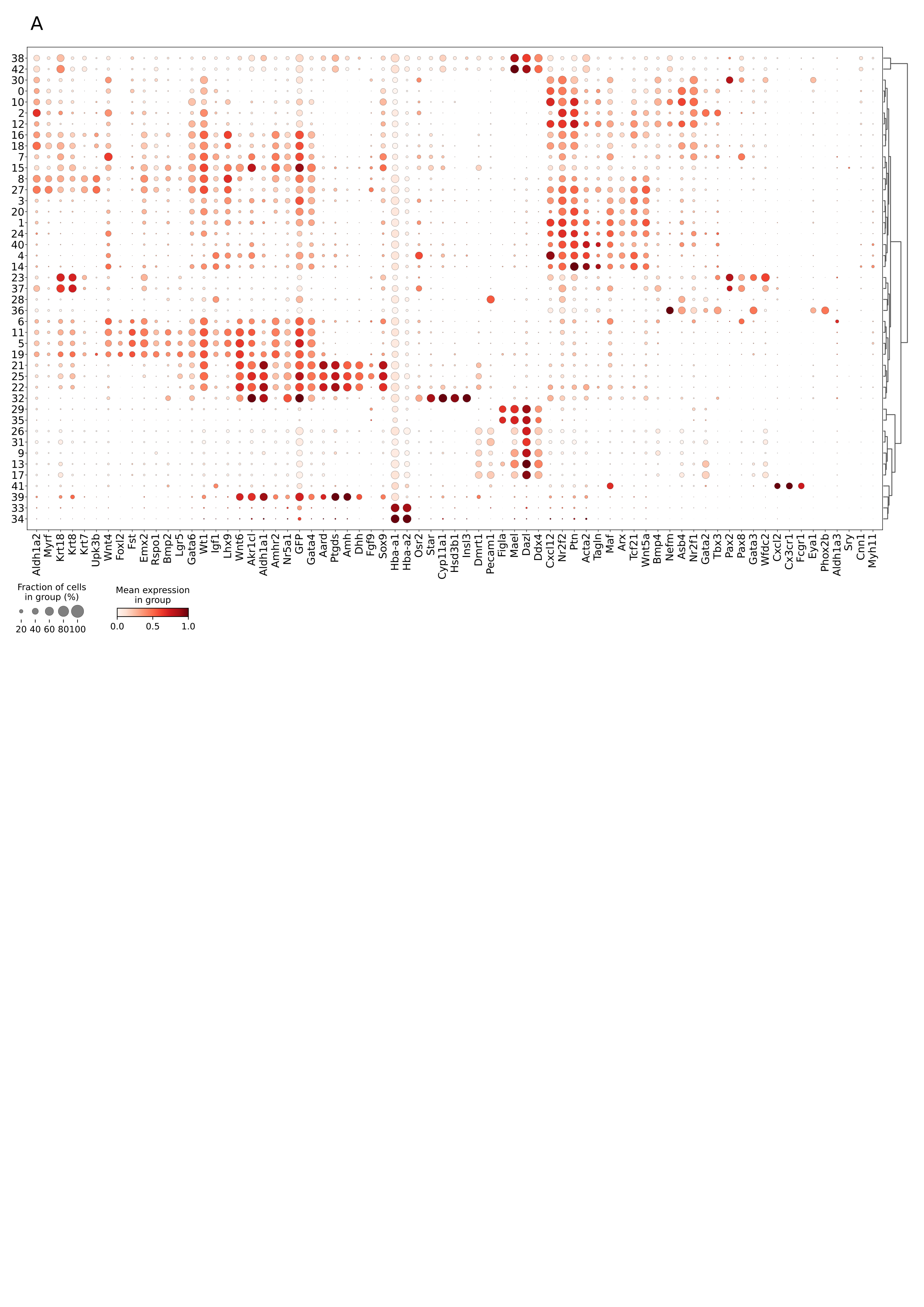

Fig. S1. Dotplot representation of the expression of marker genes in each Leiden cluster. The size of the node represents the percentage of cells in the cluster expressing the gene and the color indicates the level of expression (log normalized counts). Clusters and genes are ordered by hierarchical clustering based on Spearman correlation.

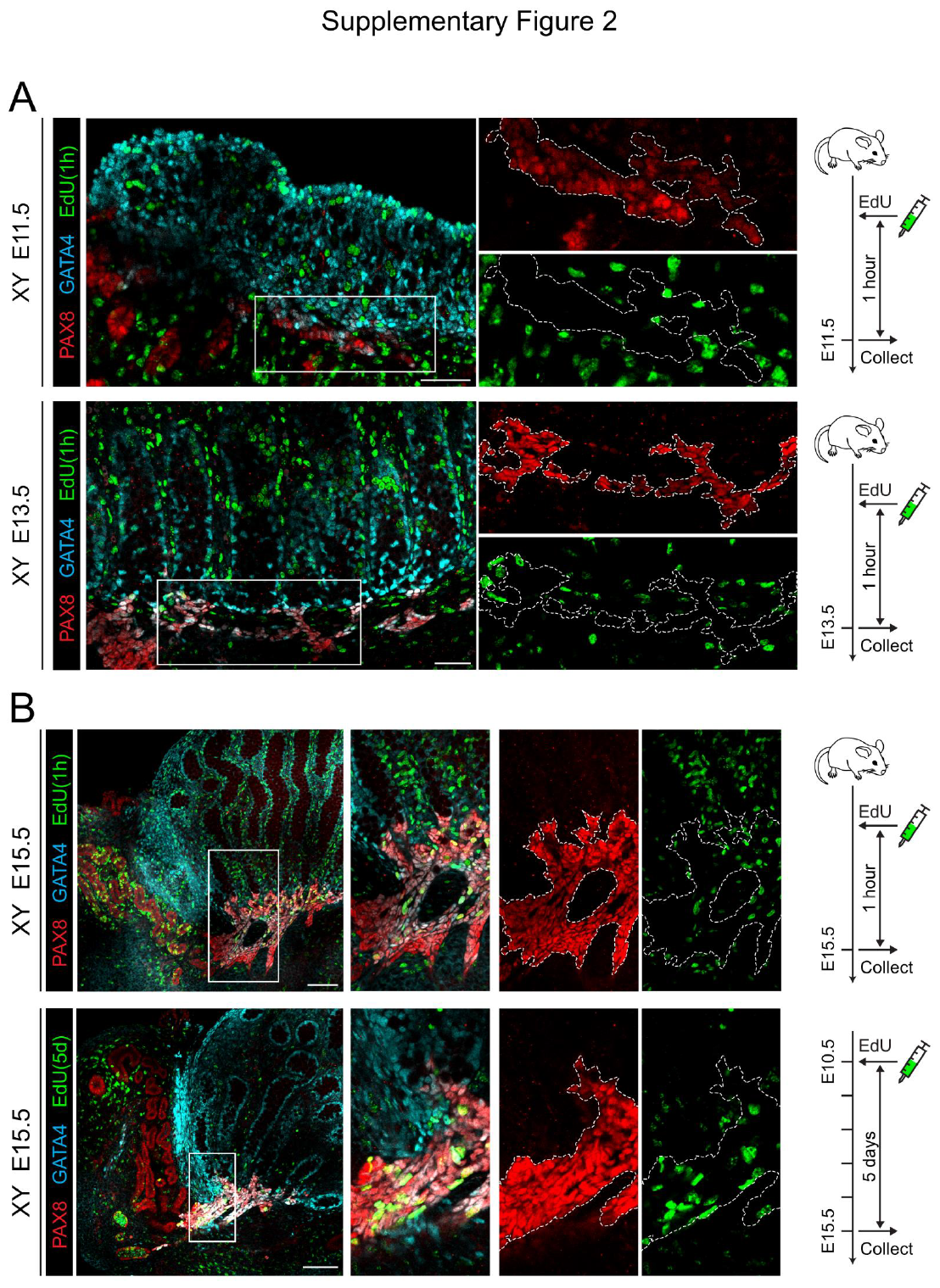

**Fig. S2.** **EdU labeling of PAX8^+^ cells reveals that these cell cycle arrest in early population and a mixed population of cells in the rete testis at E15.5.** (A-B) Wholemount immunofluorescence of gonads pulsed with EdU (green). Overlap of PAX8 (red) and GATA4 (cyan) was used to identify SLCs and the presumptive rete testis (dotted line). Boxes in merged images indicate regions shown as isolated channels on right. (A) XY gonads pulsed with EdU an hour before collection at E11.5 or E13.5. (B) XY gonads pulsed with EdU either 1 hour or 5 days before collection at E15.5. Scale bar = 50um.

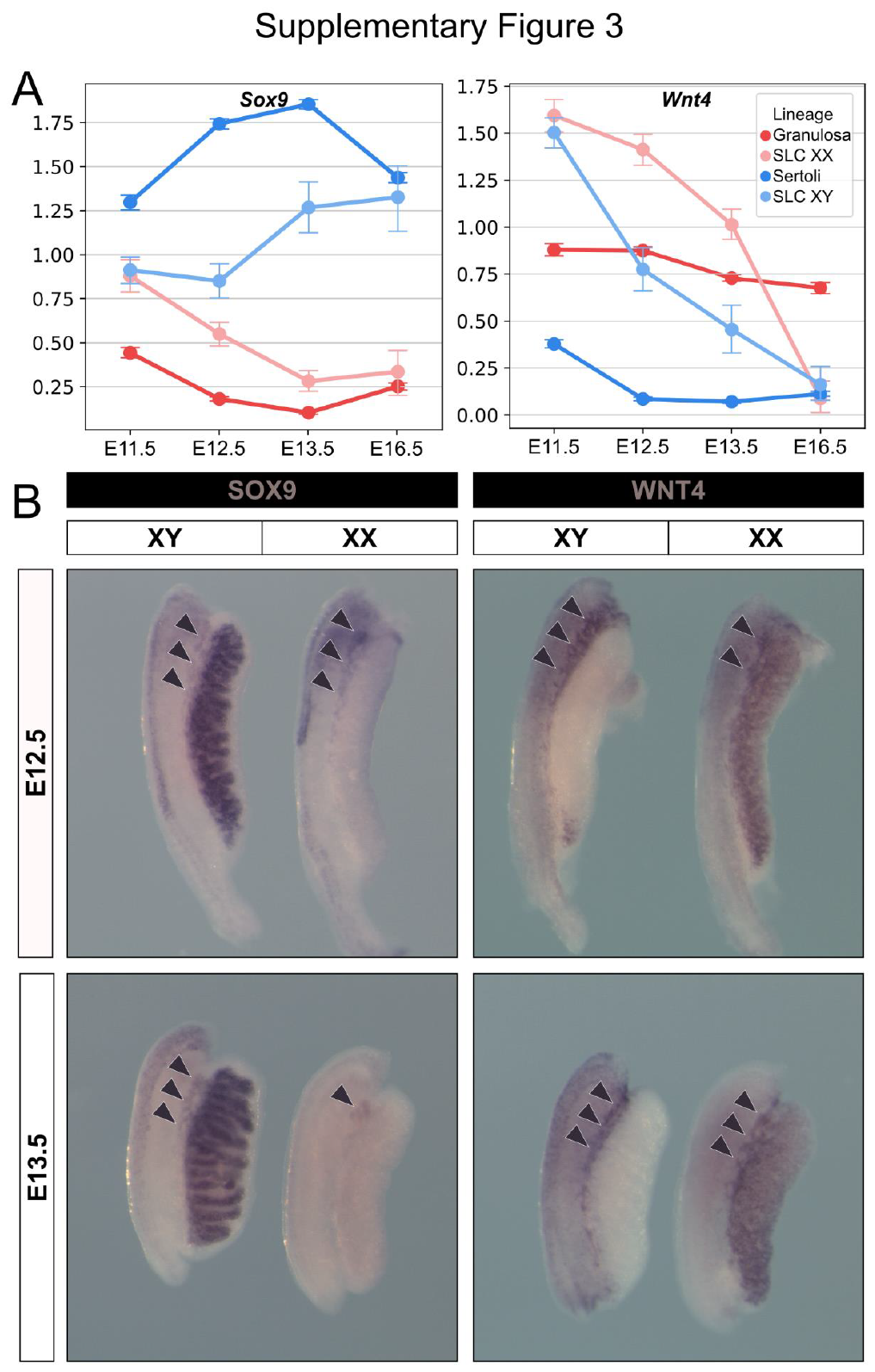

Fig. S3. *Sox9* and *Wnt4* are both express in SLC of the rete testis and rete ovarii. (A) Point plot representation of the expression of *Wnt4* and *Sox9* at the different developmental time points in both SLCs and supporting cells. Data were extracted from scRNA-seq analysis of XX and XY embryos at E11.5, E12.5, E13.5 and E16.5. (B) Whole-mount in-situ hybridization analysis of *Sox9* and *Wnt4* in XX and XY gonads at E12.5 and E13.5. Wnt4 and Sox9 transcripts were both detected in the rete testis and rete ovarii region (arrowheads).

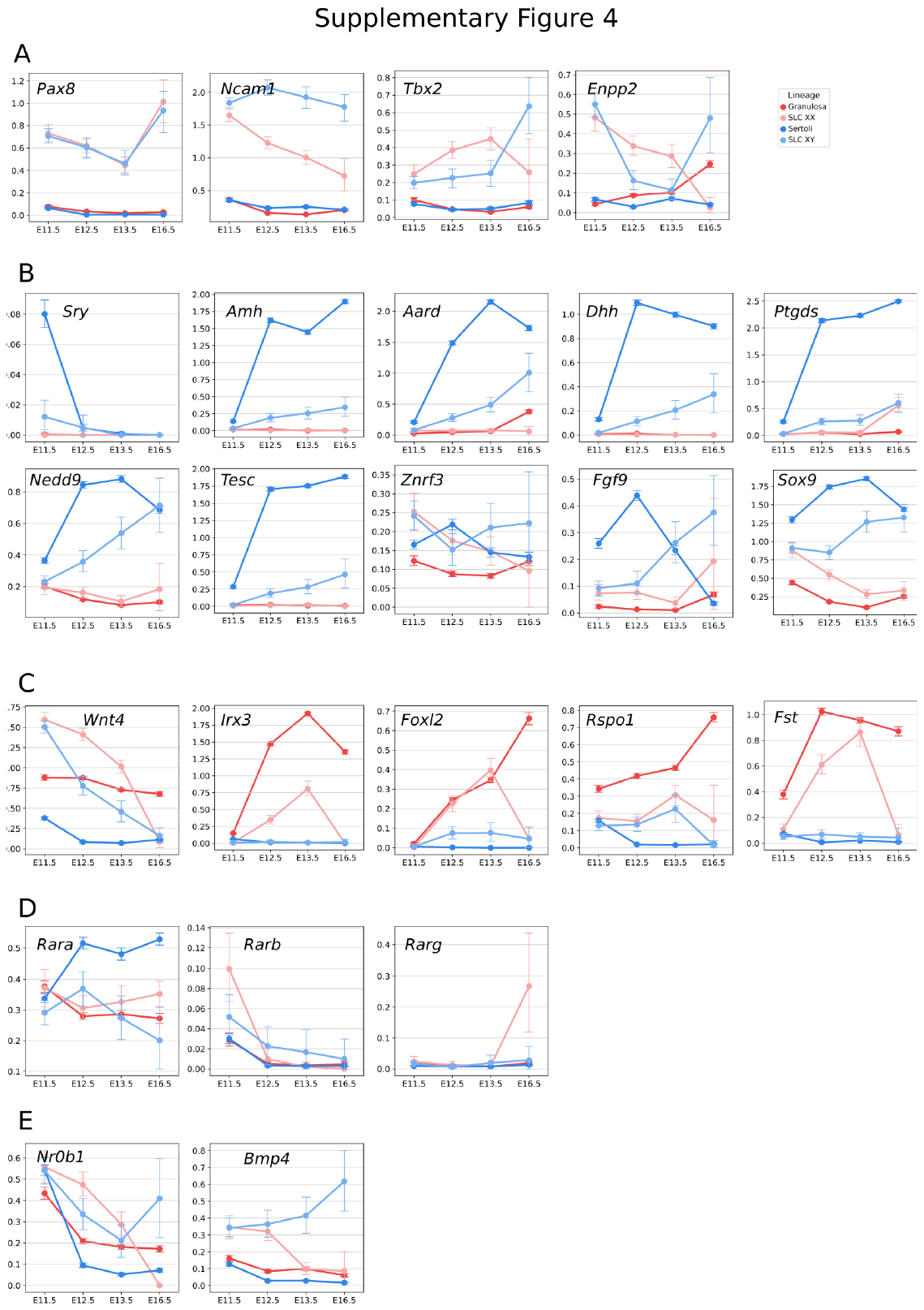

Fig. S4. Expression profiles of selected genes in SLC and supporting cells. Point plot representation of the expression of selected marker genes at the different developmental time points in both SLCs and supporting cells. Genes were classified according to their expression profiles or the relevance for a signaling pathway: (A) SLC, (B) Sertoli cells, (C) granulosa cells, (D) retinoic acid signaling and *Nr0b1* and *Bmp4* (E) miscellaneous.

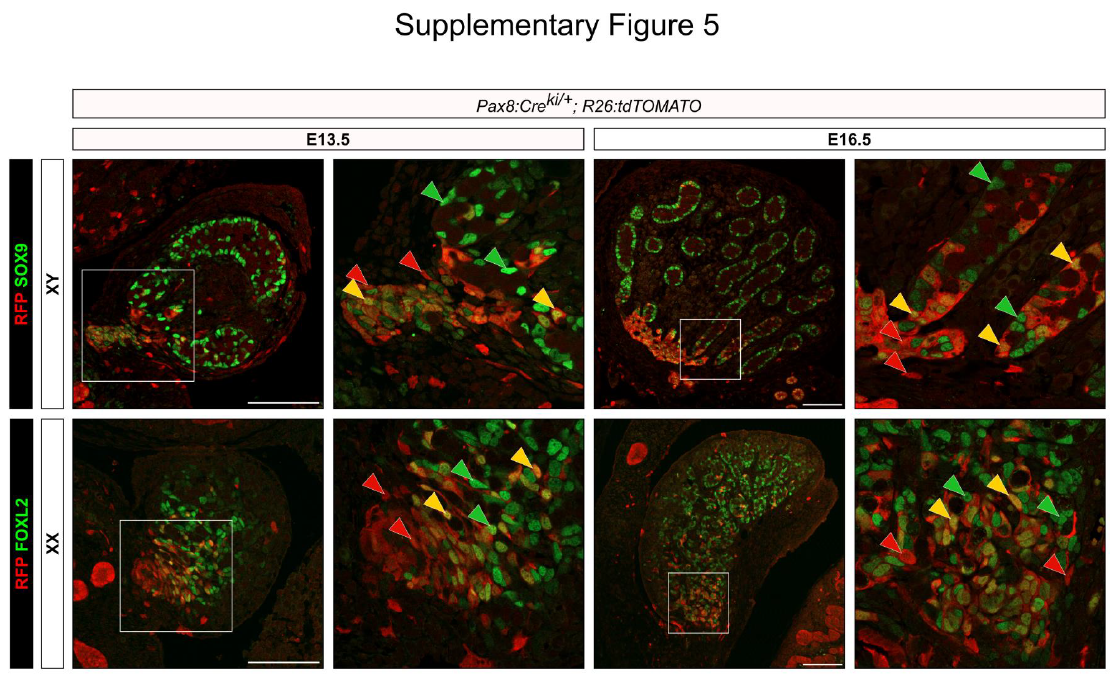

Fig. S5. SLC progenitors are multipotent and contribute to both rete structures and supporting cell lineage. Co-immunofluorescence for RFP/SOX9 and RFP/FOXL2 in XY and XX gonads of *Pax8:Cre;Rosa26:tdTomato;Nr5a1:GFP* at E13.5 and E16.5. Red arrowheads indicate RFP^+^ cells, yellow arrowheads indicate RFP^+^/SOX9^+^ or RFP^+^/FOXL2^+^ cells, green arrowheads indicate SOX9^+^ or FOXL2^+^ cells. Presence of RFP+ cells is highest near the rete. RFP^+^/SOX9^+^ cells are present in testis cords at E13.5 and E16.5. Similarly, RFP^+^/FOXL2^+^ cells are present in the ovary at E13.5 and E16.5. Scale bar 100 µm.

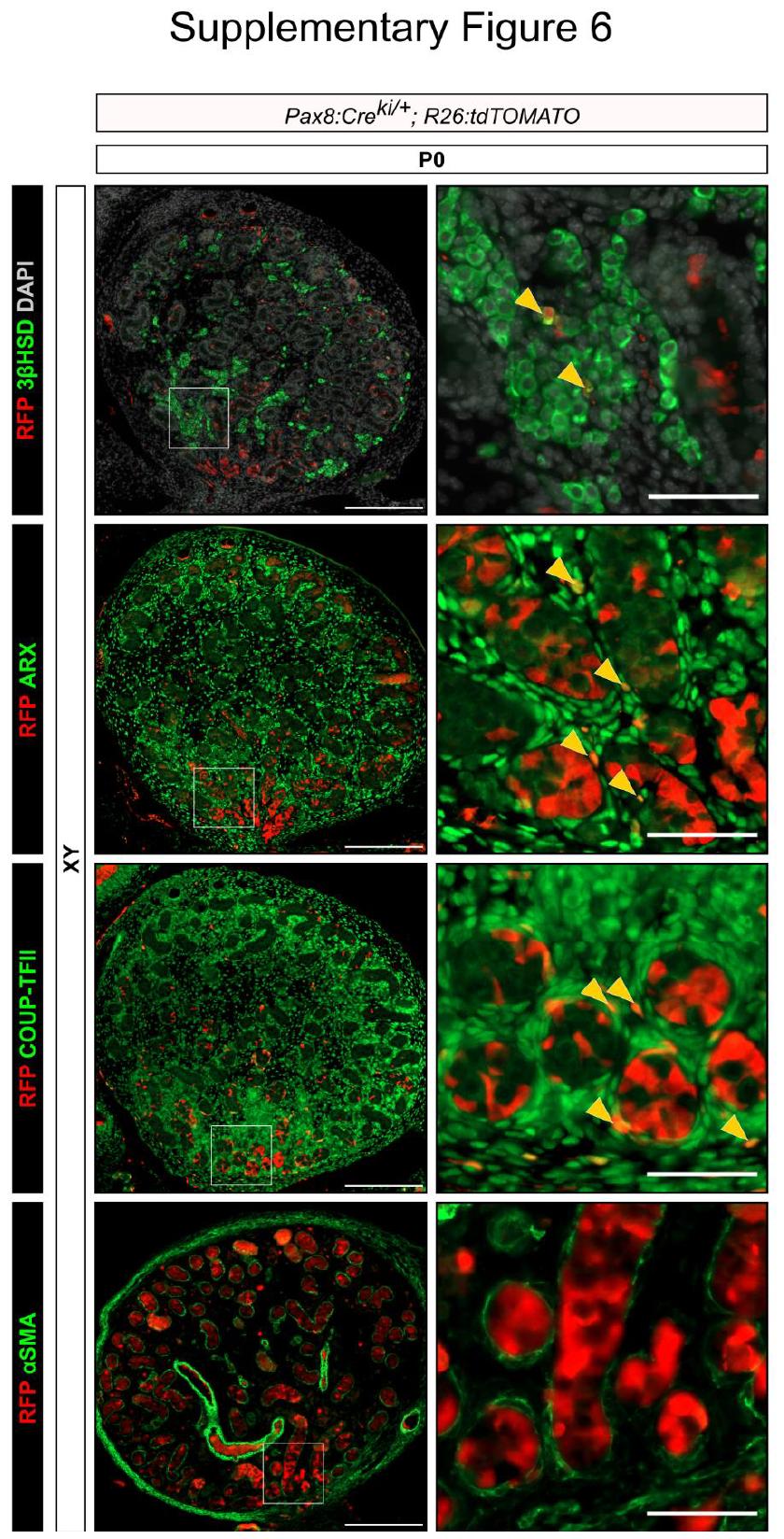

**Fig. S6. SLCs rarely contribute to the population of Leydig cells, interstitial progenitors and peritubular myoid cells in XY gonads.** Co-immunofluorescence of XY gonads of *Pax8:Cre;Rosa26:tdTomato;Nr5a1:GFP* mice at P0 for RFP and 3βHSD, a marker of the Leydig cells, RFP and ARX or COUP-TFII, two markers of interstitial progenitors, and RFP and αSMA, a marker of the peritubular myoid cells. Boxes indicate regions shown on the right. Yellow arrowheads indicate co-labeled cells. Scale bars 200 µm and 50 µm.

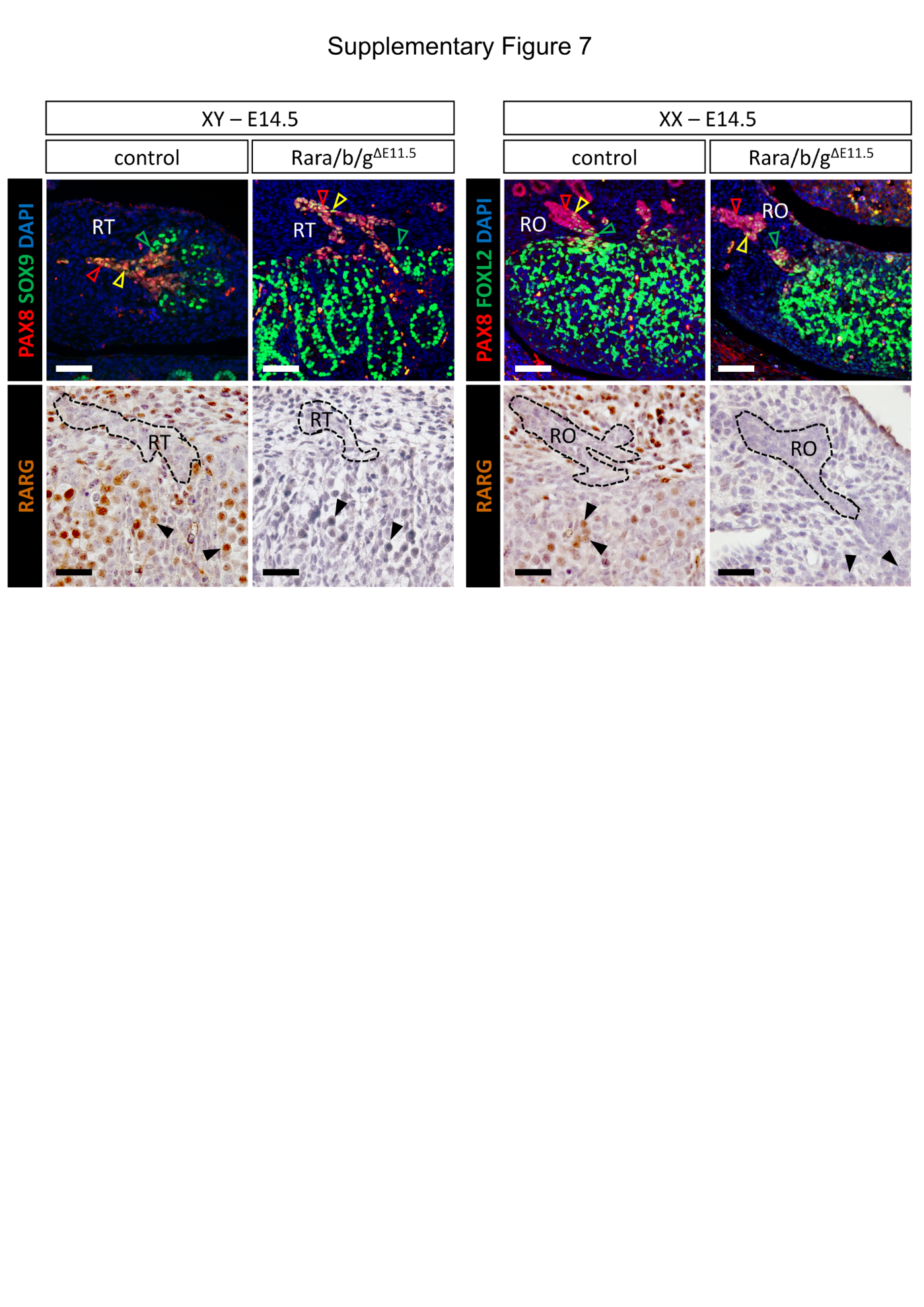

Fig. S7. Ablation of all *Rar*-coding genes from E11.5 onwards does not impair formation and differentiation of rete testis/ovarii cells. Upper left panel: immuno-detection of PAX8 (red signal), SOX9 (green signal) in control and mutant (Rara/b/g^∆E11.5^) male (XY) embryos at E14.5. Upper right panel: immuno-detection of PAX8 (red signal), FOXL2 (green signal) in control and mutant (Rara/b/g^∆E11.5^) female (XX) embryos at E14.5. Lower left panel: immuno-detection of RARG (in brown) in control and mutant (Rara/b/g^∆E11.5^) male (XY) embryos at E14.5. Nuclei were counterstained with DAPI (blue signal). Note that PAX8^+^/SOX9^+^ and PAX8^+^/FOXL2^+^ cells appear yellow. Lower right panel: immuno-detection of RARG (in brown) in control and mutant (Rara/b/g^∆E11.5^) female (XX) embryos at E14.5. Note that (i) in the control situation, RARG is detected mainly in germ cells and mesonephric mesenchymal cells, but not in rete testis/ovarii cells; (ii) in Rara/b/g^∆E11.5^ embryos, RARG is efficiently lost in virtually all cell-types, as anticipated (see (Vernet, et al., 2020)). Legend: RO, rete ovarii; RT, rete testis; XX, female fetus; XY, male fetus; the dotted lines delineate the rete testis/ovarii; red, orange and green open arrowheads point to PAX8^+^, PAX8^+^/SOX9^+^ or PAX8^+^/FOXL2^+^, and SOX9^+^ or FOXL2^+^ cells, respectively; black filled arrowheads point to RARG^+^ germ cells. Scale bars is 50 µm.

**Table S1**

| **Supplementary table 1: Relative abundance of cell types during the process of sex determination** | | | | | | | | | | |
| --- | --- | --- | --- | --- | --- | --- | --- | --- | --- | --- |
| **Developmental stage** | **E10.5** | | **E11.5** | | **E12.5** | | **E13.5** | | **E16.5** | |
| **Genetic sex** | **XX** | **XY** | **XX** | **XY** | **XX** | **XY** | **XX** | **XY** | **XX** | **XY** |
| **Cell type** |  |  |  |  |  |  |  |  |  |  |
| **Germ cells** | 0,2 | 0,8 | 5,6 | 5,8 | 27,2 | 16,6 | 27,7 | 24,3 | 34,0 | 6,0 |
| **Coelomic epithelium** | 10,4 | 20,3 | 11,3 | 17,9 | 0,1 | 0,1 | 0,0 | 0,0 | 0,0 | 0,0 |
| **Surface epithelium** | 0,0 | 0,0 | 0,2 | 0,2 | 9,0 | 12,1 | 15,9 | 7,1 | 0,6 | 2,5 |
| **Pre-supporting** | 0,0 | 0,0 | 15,3 | 16,7 | 0,0 | 0,0 | 0,0 | 0,0 | 0,0 | 0,0 |
| **Sertoli** | 0,0 | 0,0 | 0,0 | 0,6 | 0,6 | 17,1 | 0,0 | 14,4 | 0,0 | 22,1 |
| **Granulosa** | 0,0 | 0,0 | 0,1 | 0,0 | 35,9 | 0,1 | 35,1 | 0,0 | 35,4 | 0,0 |
| **Early supporting-like** | 0,0 | 0,0 | 2,7 | 2,7 | 0,0 | 0,0 | 0,0 | 0,0 | 0,0 | 0,0 |
| **Late supporting-like** | 0,0 | 0,0 | 0,0 | 0,0 | 3,6 | 1,6 | 2,0 | 0,7 | 0,1 | 0,3 |
| **Fetal Leydig** | 0,0 | 0,0 | 0,0 | 0,0 | 0,0 | 0,1 | 0,0 | 4,1 | 0,0 | 4,8 |
| **Early interst. progenitors** | 0,0 | 0,0 | 0,1 | 0,2 | 8,7 | 39,2 | 15,1 | 38,4 | 0,1 | 0,2 |
| **Late interst. progenitors** | 0,0 | 0,0 | 0,0 | 0,0 | 0,0 | 0,0 | 0,0 | 0,2 | 24,7 | 57,6 |
| **Invading Meson.** | 0,0 | 0,0 | 0,1 | 0,0 | 7,8 | 4,8 | 0,2 | 4,3 | 0,0 | 0,1 |
| **Perivascular** | 0,0 | 0,0 | 0,0 | 0,0 | 0,0 | 0,1 | 0,0 | 1,1 | 0,7 | 1,0 |
| **Mesonephric tubules** | 20,0 | 16,2 | 7,6 | 5,8 | 1,1 | 0,3 | 0,0 | 0,1 | 0,0 | 0,0 |
| **Mesonephric mesenchyme** | 64,5 | 56,7 | 54,8 | 47,3 | 0,2 | 0,9 | 0,0 | 0,0 | 0,0 | 0,0 |
| **Endothelium** | 0,7 | 1,6 | 1,2 | 1,3 | 1,9 | 4,0 | 1,3 | 2,1 | 0,6 | 0,7 |
| **Immune** | 0,3 | 0,2 | 0,3 | 0,5 | 0,3 | 0,3 | 0,2 | 0,3 | 0,5 | 0,5 |
| **Adrenosympatic** | 3,7 | 3,7 | 0,4 | 0,4 | 0,0 | 0,1 | 0,0 | 0,0 | 0,0 | 0,0 |
| **Erythrocytes** | 0,2 | 0,5 | 0,3 | 0,6 | 3,7 | 2,8 | 2,3 | 3,0 | 3,2 | 4,1 |

Captions for data S1-S4

Data S1. Differentially expressed genes between XY SLCs and supporting cells at early, late, and all stages, and associated GO biological process terms.

Data S2. Differentially expressed genes between XX SLCs and supporting cells at early, late, and all stages, and associated GO biological process terms.

Data S3. Differentially expressed genes between cells from CE sub-cluster 6 and CE sub-clusters 1 and 5.

Data S4. Sets of genes selected by LASSO modeling for granulosa and Sertoli cells and associated weights
